# Mass spectrometry reveals the chemistry of formaldehyde cross-linking in structured proteins

**DOI:** 10.1101/820779

**Authors:** Tamar Tayri-Wilk, Moriya Slavin, Joanna Zamel, Ayelet Blass, Shon Cohen, Alex Motzik, Xue Sun, Deborah E. Shalev, Oren Ram, Nir Kalisman

## Abstract

Formaldehyde is a widely used fixative in biology and medicine. The current mechanism of formaldehyde cross-linking of proteins is the formation of a methylene bridge that incorporates one carbon atom into the link. Here, we present mass spectrometry data that largely refute this mechanism. Instead, the data reveal that cross-linking of structured proteins mainly involves a reaction that incorporates two carbon atoms into the link. Under MS/MS fragmentation, the link cleaves symmetrically to yield previously unrecognized fragments carrying a modification of one carbon atom. If these characteristics are considered, then formaldehyde cross-linking is readily applicable to the structural approach of cross-linking coupled to mass spectrometry. Using a cross-linked mixture of purified proteins, a suitable analysis identifies tens of cross-links that fit well with their atomic structures. A more elaborate *in situ* cross-linking of human cells in culture identified 469 intra-protein and 90 inter-protein cross-links, which also agreed with available atomic structures. Interestingly, many of these cross-links could not be mapped onto a known structure and thus provide new structural insights. For example, two cross-links involving the protein βNAC localize its binding site on the ribosome. Also of note are cross-links of actin with several auxiliary proteins for which the structure is unknown. Based on these findings we suggest a revised chemical reaction, which has relevance to the reactivity and toxicity of formaldehyde.

## Introduction

Formaldehyde (FA) has been used as a fixative and preservative for many decades [Karnovsky, 1965; Carson, 1973]. It is reactive towards both proteins and DNA, and forms inter-molecular cross-links between macromolecules [Solomon, 1985] as well as intra-molecular chemical modifications [Chang, 1994; Metz, 2004]. The reactivity of FA in coupling with its high permeability into cells and tissues led to its use in numerous applications in biology, biotechnology, and medicine [Hoffman, 2015]. The reaction involving FA cross-linking between amino acids is assumed to involve two steps [Fraenkel-Conrat, 1948; Feldman, 1973]. In the first step, a nucleophilic group on an amino acid (typically the primary amine on lysine side chains) attacks the carbon in the FA to form a carbinol derivative, followed by a condensation reaction to form an imine (R^1^N=CH_2_). In the second step, the imine reacts with a nearby amino acid to form a methylene bridge (R^1^NH-CH_2_-R^2^), which completes the cross-linking. In terms of mass, the above reaction adds 12 Da (one carbon atom) to the total mass of the two cross-linked amino acids. Mass spectrometry has confirmed this 12 Da addition to the masses of peptides after FA incubation [Metz, 2004; Metz, 2006; Toews, 2008; Wang, 2016]. Mass spectrometry has also reported mass additions of 24, 36, 48, and 60 Da, but only after longer FA incubation times. These heavier ions were considered to be multiple occurrences of the above 12 Da (methylene bridge) reaction on the same peptide. It is important to note that the above mechanism was formulated from studies of single amino-acids, model peptides, or very small proteins (such as insulin). To the best of our knowledge, a systematic study of FA cross-linking on large and structured proteins has not been performed to date.

A potential use of FA is for the experimental technique of cross-linking coupled to mass-spectrometry (XL-MS) for structural characterization of protein assemblies [Leitner, 2016; Schneider, 2018; Sinz, 2018]. In XL-MS, a cross-linking reagent reacts with a protein complex and forms covalent links between residues that are in spatial proximity. Analysis by mass spectrometry then identifies which residue pairs underwent cross-linking. This proximity information can be used to map protein interaction networks [Herzog, 2012], and as distance restraints for structural modeling [Rappsilber, 2011]. An important advance in XL-MS is *in situ* experiments [Weisbrod, 2013; Kaake, 2014], in which intact cells or tissues are cross-linked rather than purified proteins. *In situ* XL-MS offers the advantage of reporting on the architectures of protein complexes as they occur in their native environment. Recent applications of *in situ* XL-MS were on size-reduced systems such as isolated mitochondria [Schweppe, 2017], nuclei [Fasci, 2018], and finely diced heart tissue [Chavez, 2018]. It seems that this size reduction is necessary in part because of the insufficient ability of the current cross-linking reagents to penetrate through the biological membranes.

Given its high permeability and reactivity, FA is an attractive reagent for XL-MS, and especially for *in situ* XL-MS. Yet, FA use is not common in XL-MS. We only found reports of FA being used for stabilization of protein complexes, which were later cross-linked with a different reagent for mass spectrometry identification [Robinson, 2016; Wang, 2017]. A possible cause for this puzzling situation may originate from an incorrect understanding of the FA cross-linking chemistry in structured proteins. Identification of cross-linked peptides requires accurate knowledge of the mass of the cross-link product, whereas incorrect mass will not yield any identifications. In this work, we conduct an unbiased mass-spectrometric search that indeed identifies a different reaction product for FA cross-linking with a mass of 24 Da. This new reaction only occurs in structured proteins (rather than peptides), explaining why earlier works did not observe it.

## Results

### Formaldehyde cross-linking of purified proteins

We first surveyed the FA cross-linking products that occur within structured proteins by cross-linking a mixture of three purified proteins (Bovine Serum Albumin (BSA), Ovotransferrin, and α-Amylase). The mixture was incubated with FA for twenty minutes, and then quenched, denatured, digested by a trypsin into peptides, and analyzed by mass spectrometry (Figure 1A). The general practice to identify a cross-linked peptide pair in mass spectrometry data is to match between a measured mass and the theoretical total mass of the two peptides and the cross-linker. Here, our search was not limited to one predetermined cross-linker mass, but rather scanned through a range of possible masses. Figure 1C shows the number of cross-links that the search identified for each cross-linker mass that was tested. It was surprising to see that the cross-linking is dominated by a reaction that adds exactly 24 Da (two carbon atoms) to the total mass of the two peptides. This is different from the expected 12 Da mass of the methylene bridge that was generally assumed for the FA cross-linking reaction [Fraenkel-Conrat, 1948]. The broadening of the peak, which apparently includes reactions that add 25, 26 and 27 Daltons, is a known artifact resulting from incorrect assignment of the mono-isotopic mass by the mass spectrometer (Figure S1). This artefact is common in XL-MS analysis [Lenz, 2018; Götze, 2019], and should not be interpreted as an alternative reaction. We also tested another brand of FA, which resulted in the same mass-scan profile (Figure S2).

**Figure 1.**
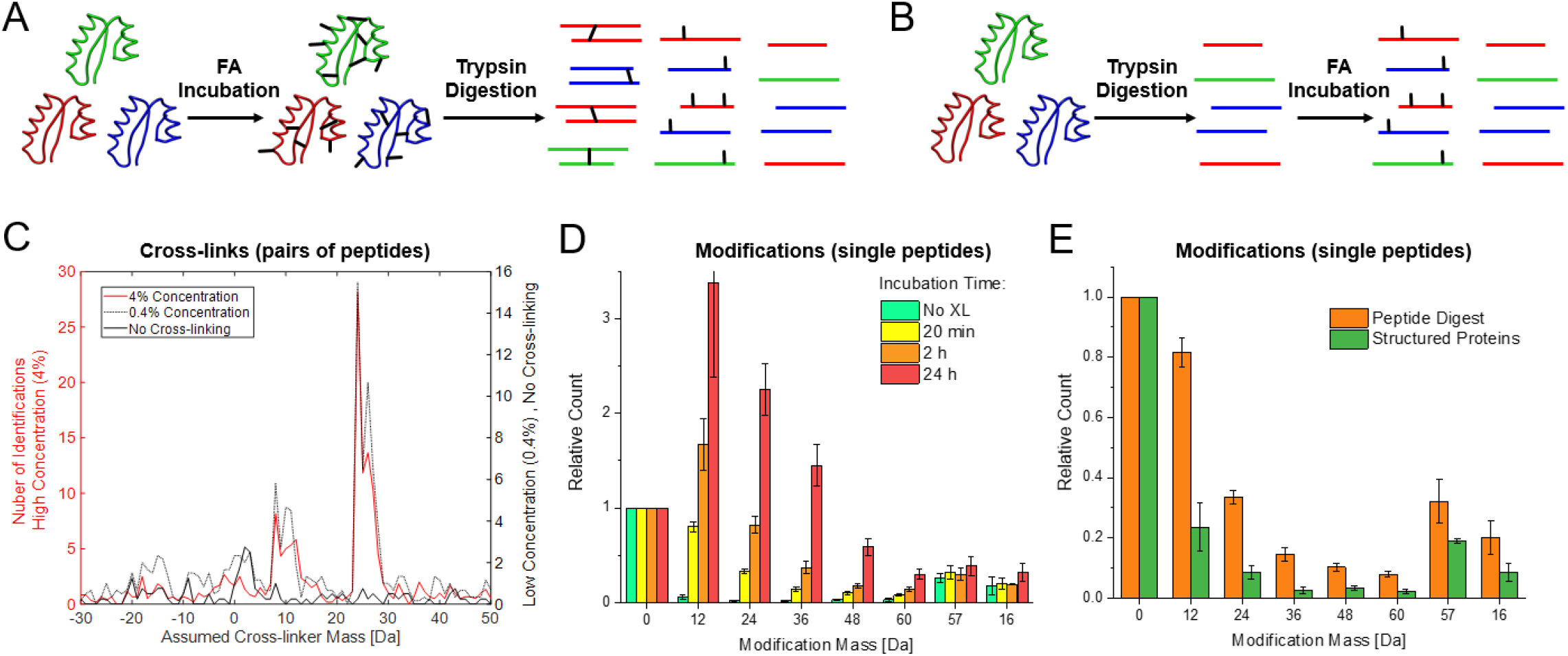
Formaldehyde (FA) cross-links are fundamentally different from local modifications. **(A and B)** Two experimental setups involving a mixture of three proteins. The final digests consist of linear peptides, peptides with FA-induced modifications, and cross-linked peptide pairs. Only setup A resulted in detectable cross-links. **(C)** Standard search for cross-links in mass spectrometry data from setup A. The Y-axis shows the number of identified cross-links. The search was repeated multiple times, each time assuming a different mass (X-axis) being added by the cross-linking reaction to the total mass of the two linked peptides. Overlaid are mass scans from three experimental conditions: high (4%) FA concentration (red), low (0.4%) FA concentration (gray), and a control without FA (black). Formaldehyde cross-linking is dominated by a reaction that adds exactly 24 Daltons to the total mass of the two cross-linked peptides. **(D)** Search for FA modifications on single peptides in mass spectrometry data from setup B. Each bar series is normalized by the count of non-modified peptides (‘0’ modification mass). We searched for matches to multiple occurrences of the 12 Da reaction on the same peptide (24, 36, 48, and 60 Da). To give scale to these results, we also searched for peptides with off-target alkylation (57 Da) and peptides with oxidized methionine residues (16 Da). For all FA incubation times, we see that peptides with a single 12 Da modification are the most frequent. **(E)** A similar search for modifications, but this time comparing setup B (‘peptide digest’) to setup A (‘structured proteins’). In both setups, the FA concertation was 2% and the incubation time was 20 minutes. Formaldehyde is less reactive towards structured proteins, forming fewer modifications.

The 24 Da reaction is not two separated 12 Da reactions occurring in parallel within the same peptide pair for two reasons: First, one expects that a lower concentration of FA will show less of the 24 Da reaction and more of the 12 Da reaction. Yet, for both high and low concentrations (Figure 1C) the mass-scan profiles are the same, albeit there are less identifications at the lower concentration. Second, ion species corresponding to mass additions of 36 Da or 48 Da are not observed, further strengthening the argument that parallel links are not the underlying cause.

As a control to the experiments on structured proteins, we incubated the peptide digest from the three-protein mixture with FA, and analyzed the products by mass spectrometry (Figure 1B). This analysis did not identify any cross-link that formed between peptide pairs of the digest. Yet, an analysis of single linear peptides found a high abundance of FA-related modifications (Figure 1D). Even after 20 minutes of incubation with 2% FA, the number of peptides with a 12 Da modification was nearly equal to that of peptides without modifications. These modifications were nearly absent when the digest was not treated with FA (‘no XL’), and can therefore be attributed to the FA reactivity. Peptides with multiple modifications in parallel were also frequent, and became more so at longer incubation times. The experimental setup in Figure 1A also led to formation of linear peptides with FA modifications (Figure 1E). Yet, fewer modifications are observed when FA is incubated with structured proteins compared with digest. This may be due to the shielding of modification sites in the core of the protein structure. Overall, the data show that chemistry of local modifications is different from that of long-range cross-linking. Whereas a 12 Da reaction is the most prevalent for the former, a 24 Da reaction is by far dominating the latter.

Further support for the uniqueness of the 24 Da reaction is seen in the unusual fragmentation pattern of its MS/MS spectra (Figure 2A-C). The cross-link is highly susceptible to higher-energy collisional dissociation (HCD), and fragments in which it stayed intact could not be detected. Instead, it breaks symmetrically to give a mass addition of 12 Da on each peptide. Peaks corresponding to the total mass of one of the peptides plus 12 Da were among the most intense in the observed MS/MS spectra. The two peptides then break a second time to yield the standard b- and y-fragments as well as modified b- and y-fragments with an additional 12 Da mass. The unique fragmentation pattern associated with the 24 Da reaction bears resemblance to that of cleavable cross-linkers that are now frequently used in XL-MS [Sinz, 2017]. Yet, an important distinction is the 100% cleavage efficiency of the 24 Da reaction, which is higher than that observed with other cleavable cross-linking reagents. This unique fragmentation may be another cause to why the 24 Da reaction was not reported before in mass spectrometry applications.

**Figure 2.**
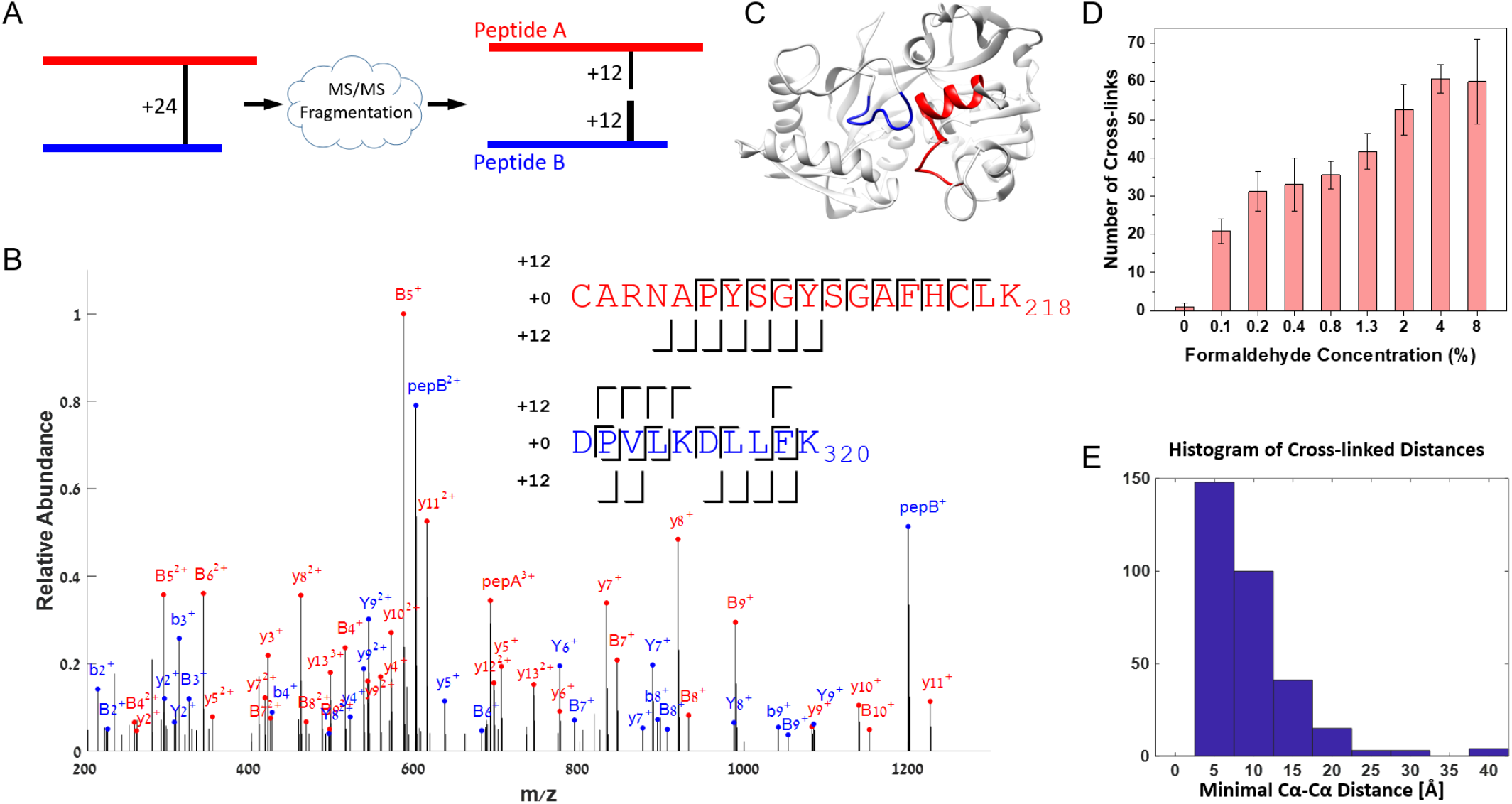
**(A)** The MS/MS fragmentation of a formaldehyde cross-link breaks the ion to its two peptides, each with an additional mass of exactly 12 Da. The two peptides then break a second time to produce b- and y-fragments. **(B)** The MS/MS spectrum of a cross-link between two peptides of Ovotransferrin. PepA and PepB are peaks matching the total mass of the corresponding peptides plus 12 Da. Peaks annotated with capital ‘B’ or ‘Y’ match the masses of the corresponding b- and y-fragments plus 12 Da. For clarity, only peaks with intensities in the top quartile are shown. The annotation of the full spectrum is shown in Figure S3. **(C)** The two peptides identified in the above MS/MS spectrum are spatially close to each other in the crystallographic structure of Ovotransferrin. **(D)** The average number of cross-links identified for each formaldehyde concentration on a mixture of three proteins. Standard deviation calculated across six replicates. The estimated false detection rate is 3%. **(E)** Histogram of the distances between cross-linked peptides identified in the 4% formaldehyde samples. The cross-link distance is estimated by the minimal Cα-Cα distance between the peptide pair on the crystallographic structure.

With the understanding of the unique properties of FA XL-MS, we designed an analysis application that is different from the one used for the mass scans. The new application is tailored specifically to identify the 24 Da reaction and its subsequent MS/MS pattern with the modified plus-12 Da fragments. The application successfully identified cross-links in the three-protein mixture in a concentration-dependent manner (Figure 2D). Interestingly, the application could also detect a small number of cross-links corresponding to the 12 Da reaction, but at a ratio of less than 1:7 compared with the 24 Da reaction. The three proteins did not cross-link equally: most of the identifications were in BSA and Ovotransferrin, with only 2-3 identifications in α-Amylase. Non-native cross-linking between different proteins (such as BSA to Ovotransferrin) was negligible, and occurred only for FA concentrations of 4% and 8% at a frequency of less than one identification per mass spectrometry run. Table S1 lists an example of the identifications from one cross-linking experiment. An attempt to analyze the same data with an application tailored for cleavable cross-linkers [Iacobucci, 2018] gave only a third of the identifications (listed in Table S2), which were a subset of our results. The smaller number is caused by certain features of FA cross-linking, such as multiple link sites, that are currently not supported by the latter application.

The fragmentation pattern of the 24 Da reaction does not enable identification of the exact two residues that underwent cross-linking. As a typical example, the fragmentation pattern of the peptide pair shown in Figure 2B is consistent with the cross-link occurring on any of the first four residues in the upper (red) peptide. The localization is also ambiguous in the lower (blue) peptide as the first aspartic residue and the middle lysine-aspartic residues are all likely sites for the cross-link given the fragmentation. Therefore, the MS measurement shown in Figure 2B may actually report a group of isomers of the same two peptides with different cross-link sites on each. This ambiguity is unlike the fragmentation patterns encountered with cross-linking reagents that have high chemical specificity towards one amino acid type. With the latter, precise identification of the two cross-linked sites is usually possible. The uncertainty in localizing the cross-link sites prevents the measurement of the exact distance that is spanned by a FA cross-link. Instead, we estimated the cross-link distance as the minimal Cα- Cα distance between the two peptides on the protein structure. Figure 2E shows the histogram of the minimal distances observed for the cross-links from the 4% FA samples. The median distance is 8 Å, but the histogram right tail extends to values of 25-30 Å. These are typical values in XL-MS, which indicate that the cross-links fit well to the crystallographic structures of the three proteins. Figure S4 shows the histograms of the other FA concentrations and results from experiments with the common reagent disuccinimidyl suberate (DSS). This comparison indicates that FA cross-links are shorter than those of DSS.

### *In situ* formaldehyde cross-linking of human cell culture

We employed our understanding of the 24 Da reaction to identify FA cross-links from *in situ* cross-linking experiments of intact human cells. PC9 adenocarcinoma cells were incubated in 1%, 2%, 3%, 4.5%, or 6% FA solutions for 10 minutes. After the FA was washed out, the cells were lysed and the protein content prepared for mass spectrometry [Wiśniewski, 2009]. We measured 10% of the peptide digest from each FA concentration directly in the mass spectrometer. The other 90% were enriched for cross-linked peptides using strong cation exchange [Klykov, 2018], and then measured in the mass spectrometer. Standard proteomics analysis identified in the digests a set of 1692 proteins with medium to high abundance. The analysis application that has been used for the three-protein mixture could not process data of such complexity, and a different strategy was required. To that end, we took advantage of the complete dissociation of the FA cross-link during HCD, which allows matching each peptide in the pair to the data independently of the other. An application implementing this strategy analyzed each mass spectrometry run against the database of 1692 proteins in about five minutes (Methods).

Overall, the *in situ* cross-linking experiments resulted in 59 data-dependent mass spectrometry runs. The analyses of these runs searched for two separate cross-linker masses: 12 and 24 Da. We then pooled together all the identifications from these analyses into a non-redundant list of 559 cross-links (Table S3). A cross-link was reported in that list if it fulfilled the following criteria: (i) the total theoretical mass of the two peptides plus the cross-linker matched the measured precursor mass to 4 ppm; (ii) each of the two peptides is at least 8 residues long; (iii) each peptide has at least 19 fragments (of types b, y, B, and Y) matching to the MS/MS spectrum, or the ratio of the number of matching fragments to its length was higher than 1.8; and (iv) there was no other peptide pair or linear peptide that matched the MS/MS event with better fragmentation. If a cross-link occurred in multiple runs, then the occurrence with the best fragmentation was reported. The above thresholds are set to impose a false detection rate (FDR) of 3% on the entries in the list, based on decoy analysis that spiked the search database with reverse sequences.

In the list of *in situ* cross-links, the 24 and 12 Da cross-linking reactions accounted for 74% and 26% of the cross-links, respectively. This reaffirms the dominance of the 24 Da reaction in FA cross-linking also in the case of in-situ FA applications. Interestingly, the 12 Da reaction is more prevalent *in situ* than it was on the mixture of purified proteins. The identified cross-links occur within a subset of 276 proteins that are of relatively high abundance in the PC9 cell line [Geiger, 2012]. This is expected because we did not enrich for any particular protein. Encouragingly, these proteins originate from the nucleus (histones), cytoplasm (ribosomes and TRiC/CCT), mitochondria (HSP60), and endoplasmic reticulum (BiP). Such variety indicates that the FA reached most cellular compartments. We could map 280 of the cross-links onto solved atomic structures that included their corresponding peptides. Figure 3A shows the histogram of the minimal Cα- Cα distances spanned by these cross-links. The histogram includes only cross-links for which the two peptides are not consecutive in the protein sequence. The FA cross-links fit well with the atomic structures, having a minimal Cα- Cα distance below 25 Å for 272 of them (97%).

**Figure 3.**
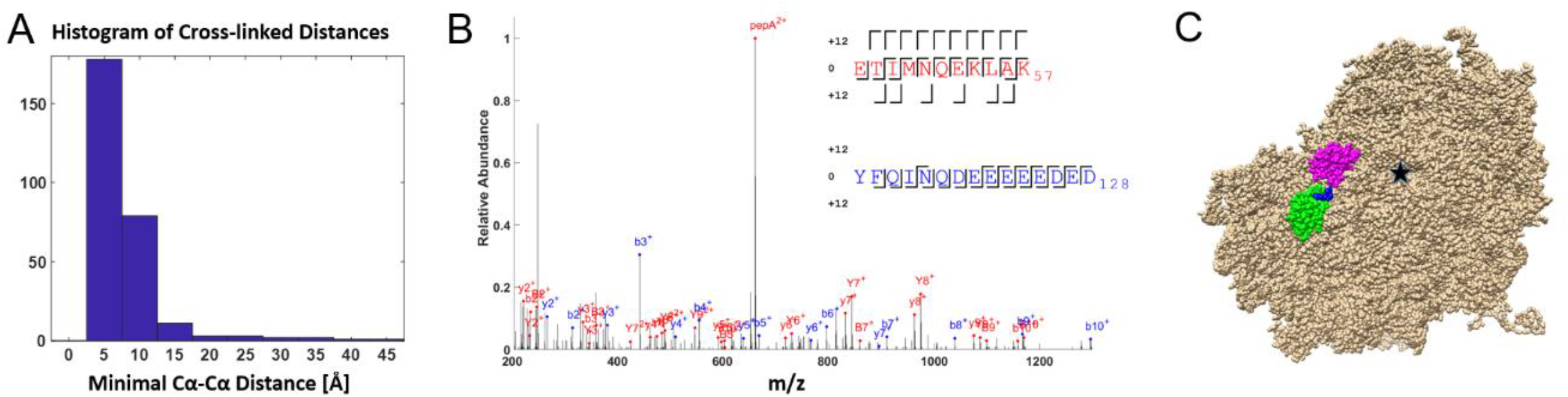
**(A)** Histogram of the distances spanned by *in situ* cross-links from cultured human cells. **(B)** Annotated MS/MS spectrum of one of the cross-links between βNAC (red peptide) and ribosomal protein L22 (blue peptide). **(C)** The outer surface of the S80 ribosome (PDBid 6ek0; Natchiar, 2017) centered on the peptide exit tunnel (black star). L22 is marked in green and its cross-linked peptide in blue. Ribosomal protein L31, which was also shown to interact with βNAC [Pech, 2010], is marked in magenta. The two cross-links found in this study suggest that βNAC binds on the ribosome in the cleft between L22 and L31.

Of the 559 cross-links, 90 (16%) are inter-protein (between two different proteins in a complex) and the rest are intra-protein (within the same protein polypeptide). A subset of 28 inter-protein cross-links had no corresponding atomic structures. Yet, they showed strong indications of being true positives. All had good fragmentation on both peptides (20 fragments or more on the weakest peptide), and most were previously reported to be part of a protein complex (Table 1). These cross-links provide new structural data - of *in situ* origin - on the relevant interactions. Particularly, each cross-link narrows down the interaction site to the vicinity of the two peptides. This information was previously unknown for most of the entries in Table 1, even when the interaction is well established. We highlight two of the cross-links that report on an interaction between the nascent polypeptide-associated (NAC) complex and the ribosome (Figure 3B). Human NAC is a cytosolic heterodimer that contacts the nascent polypeptide chain emerging from the ribosome. Previously, Pech *et al*. (2010) showed that a conserved sequence region in βNAC, which in humans spans from Thr_48_ to Leu_58_, is the interacting sequence with the ribosome. Moreover, they demonstrated by *in-vitro* cross-linking that βNAC binds to ribosomal protein L31. Here, two *in situ* cross-links include this sequence region, and couple it to the C-terminal of ribosomal protein L22. L22 and L31 are adjacent proteins on the outer surface of the ribosome (Figure 3C). Thus, the *in situ* cross-links provide further evidence to the findings of Pech *et al*., and suggest that the most likely binding site of βNAC to the ribosome is the cleft between L22 and L31.

**Table 1.**
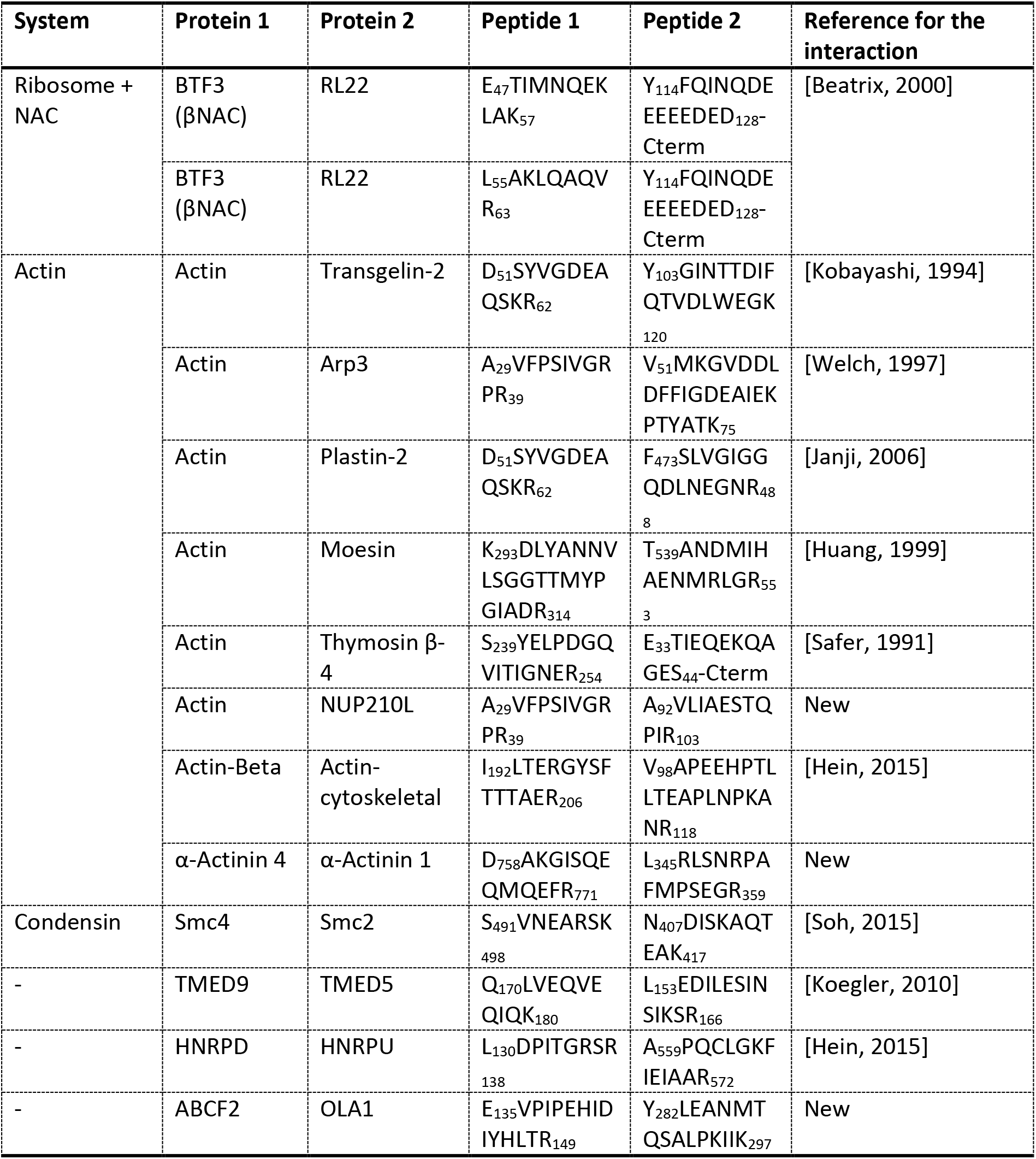
Some of the high confidence inter-protein cross-links that provide new *in situ* structural information. Annotated MS/MS spectra corresponding to these cross-links are compiled in Figure S5.

In contrast to the cross-links in Table 1, a subset of nine inter-protein cross-links had two markings of being false positives. First, they had marginal fragmentation values on one of the peptides (14-19 fragments). Second, the two cross-linked proteins were never reported to be interacting. Assuming that all the intra-protein cross-links are correct, then these cross-links comprise the false positive subset of the entire list. Being 1.6% of the list (9 out of 559), they are in accord with our *a priori* estimation of the FDR.

## Discussion

We established four features of long-range FA cross-links in proteins: First, they occur only in structured proteins. Hence, the reliance of many previous studies on peptide assays incorrectly classified the prevalent 12 Da modification as a cross-link. Second, the main cross-linking reaction involves two carbon atoms and not one. Third, these links are very labile and cleave completely under MS/MS fragmentation. Finally, the most intense MS/MS fragmentation products carry an unusual 12 Da modification. We think that all these factors contributed to the fact that the chemistry of the long-range FA cross-link has not been characterized correctly.

In light of the new findings, we suggest a new mechanism of FA cross-linking (Figure 4). The reaction starts with the accepted imine formation on the side chains of lysines. The imine formation is in accord with the prevalent 12 Da modification that others and we have observed on peptides and proteins. However, the cross-link itself forms by a dimeric interaction of two imines [Layer, 1963]. This symmetric formation is compatible with two observations. First, it explains the symmetrical cleavage of the link under MS/MS fragmentation. Second, if one assumes that the imine modification is only mildly reactive, then it is clear why cross-linking is occurring only in structured proteins: The stable structure of the protein keeps the modifications in proximity for sufficient time for the linking to occur. The mild reactivity of the imine modifications also suggests that they underlie the acute toxic effects of formaldehyde, rather than protein cross-linking.

**Figure 4.**
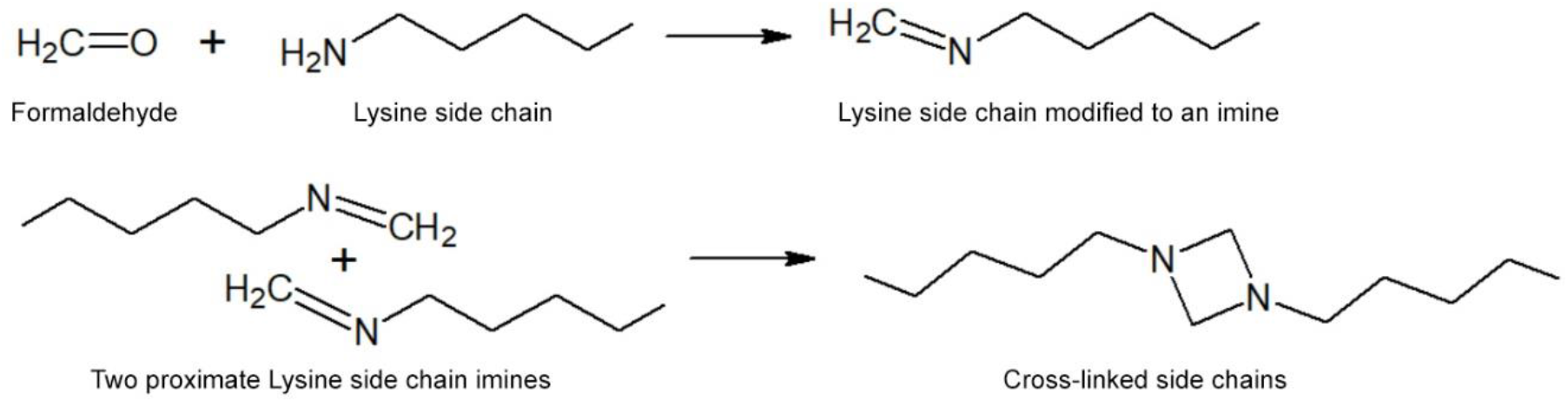
Proposed mechanism for the 24 Da reaction, demonstrated on two lysine side chains. In the first step, formaldehyde modifies the primary amine on the lysine side chain into an imine. In the second step, two modified side chains dimerize to form the cross-link product. The second step is presumably slow and can occur only in structured proteins, in which the two side chains are close and nearly stationary relative to each other.

In Figure 4 the chemical structure is exemplified on two lysine side chains, but FA cross-linking does not necessarily require two lysines. Indeed, for many of the *in situ* cross-links (Table S3) one of the peptides has no lysine residue. Therefore, the hypothesized model would have to be revised for cross-linking in the more general case. At this stage we do not have data, other than that of mass spectrometry, to support this hypothesis. So far, our attempts to support it by NMR were not successful due to the low yield of homogenous cross-linking reaction products.

## Data Deposition

The mass spectrometry data have been deposited to the ProteomeXchange Consortium via the PRIDE [Perez-Riverol, 2019] partner repository with the dataset identifier PXD015435.

## Acknowledgments

This work was supported by the Israel Science Foundation grant number 1768/15. We thank Uri Raviv for his help and advice in various stages of this work. We thank David Morgenstern and Dina Schneidman for critical reading of the manuscript.

## Methods

### Cross-linking of the three-protein mixture

A mixture solution of three purified proteins was prepared by reconstituting lyophilized protein powder in PBS (all reagents were purchased from Sigma unless noted otherwise). The proteins were Bovine Serum Albumin (BSA), Ovotransferrin, and α-Amylase with respective final molarity in the mixture of 10 μM, 10 μM, and 20 μM. Each cross-linking experiment occurred in 108 μL of solution comprising a total protein mass of 260 μg. In most experiments we cross-linked with a formalin solution (37% formaldehyde and 10% methanol) from Sigma (product number F8775). We also tested formalin with the same composition from another brand (DAEJUNG chemicals, Korea, product number 4044-4400). The formalin was incubated with the protein mixture at the desired formaldehyde concentration and the cross-linking reaction occurred at room temperature under gentle agitation. The cross-linking incubation time was 20 minutes. The cross-linking reaction was quenched by addition of ammonium bicarbonate to a final concentration of 0.5 M for 10 minutes before proceeding to mass spectrometry preparation. The results of each experimental condition are an average of six mass spectrometry runs from three experimental replicates, each with two technical replicates.

### Cross-linking of digest from the three-protein mixture

Peptide digest was prepared from the three-protein mixture by trypsin digestion as described in the ‘Mass spectrometry’ subsection ahead. The peptides were desalted on SepPak C18 column, eluted, dried in SpeedVac, and reconstituted in PBS. Formaldehyde was added to a concentration of 2% and the incubation time was either 20 minutes, 2 hours, or 24 hours. The solution was quenched by addition of ammonium bicarbonate to a final concentration of 0.5 M for 10 minutes. The peptides were desalted on C18 stage-tips and eluted for mass spectrometry analysis. The results of each incubation time are an average of two experimental replicates, each with two technical replicates.

### *In situ* cross-linking of PC9 cells

Human lung cancer cell line PC9 were seeded in Dulbecco’s modified Eagle’s medium (DMEM) (Sigma), and were supplemented with 1X Penicillin-Streptomycin (Gibco Invitrogen) and 10% fetal bovine serum (Biological Industries) at 37 °C under 5% CO_2_/95% air. The cells were grown to 80% confluency in 10 cm plates. The growth media was removed and the cells washed three times with 3 mL of warm PBS buffer. We added to each plate 2 ml of PBS with FA at different concentrations: 1%, 2%, 3%, 4.5% or 6%. The cells were incubated with FA for 15 minutes at 37°C, and then washed three times with cold PBS to remove the formaldehyde. We incubated the cells with hypertonic buffer (50 mM HEPES pH=7.5, 500 mM NaCl, 0.5 mM EDTA, 0.0005% Tween20) for 15 minutes, and then scraped the cells from the plate. The cells were centrifuged at 4°C and the supernatant was discarded. The cell pellet was resuspended for 15 minutes with hypotonic buffer (above buffer without NaCl), and then further lysed with sonication (5 seconds on, 5 seconds off, 5 times). The cell lysate was centrifuged at 4°C and the supernatant was collected. The lysate was processed by the filter-aided sample preparation (FASP) protocol [Wiśniewski, 2009] in order to remove the detergent and nucleic acids prior to the mass spectrometry analysis.

### Enrichment of cross-linked peptides by strong cation exchange chromatography (SCX)

We followed the SCX protocol by Klykov *et al*. (2018). Briefly, desalted peptide digest was dried in SpeedVac and reconstituted in 50 μl of buffer A (20% Acetonitrile, formic acid titrated to pH of 3.0). Separation was performed with an Äkta Pure system on a 100×1.0 mm PolySULFOETHYL A SCX column (PolyLC, USA) using a gradient of buffer B (20% Acetonitrile, 0.5 M NaCl, formic acid titrated to pH of 3.0) and 100 μl fractions. Fraction corresponding to NaCl concentrations of 100 mM and higher were desalted and used for mass spectrometry analysis.

### Mass spectrometry

The proteins were precipitated in acetone at −80 °C for one hour followed by centrifugation at 10,000 g. The pellet was resuspended in 20 μl of 8 M urea with 10 mM DTT. After 30 minutes, iodoacetamide was added to a final concentration of 50 mM and the alkylation reaction proceeded for 30 minutes. The urea was diluted by adding 200 μl of digestion buffer (25 mM TRIS pH=8.0; 10% Acetonitrile), trypsin (Promega) was added at a 1:100 protease-to-protein ratio, and the protein was digested overnight at 37 °C under agitation. Following digestion, the peptides were desalted on C18 stage-tips and eluted by 55% acetonitrile. The eluted peptides were dried in a SpeedVac, reconstituted in 0.1% formic acid, and measured in the mass spectrometer. The samples were analyzed by a 120 minute 0-40% acetonitrile gradient on a liquid chromatography system coupled to a Q-Exactive Plus mass spectrometer (Thermo). We were careful not to raise the temperature of the sample above 40 °C through all the preparation stages (alkylation, digestion, desalting, and in the analytical column of the LC) in order not to break the formaldehyde cross-links. The RAW data files from the mass spectrometer were converted to MGF format by Proteome Discoverer (Thermo), which was the input format for our analysis applications. The method parameters of the run were: Data-Dependent Acquisition; Full MS resolution 70,000; MS1 AGC target 1e6; MS1 Maximum IT 200 ms; Scan range 450 to 1800; dd-MS/MS resolution 35,000; MS/MS AGC target 2e5; MS2 Maximum IT 300 ms; Loop count Top 12; Isolation window 1.1; Fixed first mass 130; MS2 Minimum AGC target 800; Charge exclusion: unassigned,1,2,3,8,>8; Peptide match - off; Exclude isotope - on; Dynamic exclusion 45 seconds.

### Scanning for the mass of the cross-linking reaction (Figure 1C)

We modified our analysis application, FindXL [Kalisman, 2012], so that it ran multiple times, each time with a different cross-linker mass. We scanned all the integer masses from −30 to 50 Da. FindXL exhaustively enumerates all the possible peptide pairs and compare them to the measured MS/MS events in search of matches that fulfill the criteria below. The search parameters were: Sequence database – the sequences of BSA, Ovotransferrin, and α-Amylase; Protease – trypsin, allowing up to three miscleavage sites; Fixed modification of cysteine by iodoacetamide; Variable modification of methionine by oxidation; Cross-linking can occur on any residue type; Cross-linker is non-cleavable; MS/MS fragments to consider – b-ions and y-ions as well as b-ions and y-ions with the additional mass of the second peptide and the cross-linker; MS^1^ tolerance – 6 ppm; MS^2^ tolerance – 8 ppm.

A cross-link was identified as a match between a MS/MS event and a peptide pair if it fulfilled four conditions: 1) The mass of the precursor ion is the same as the expected mass of the cross-linked peptide pair within the MS^1^ tolerance; 2) At least four MS/MS fragments (within the MS^2^ tolerance) were identified on each peptide; 3) The fragmentation score of the cross-link (defined as the number of matching MS/MS fragments divided by the combined length of the two peptides) is 0.6 or higher; 4) The peptides are not overlapping nor consecutive in the protein sequence. The purpose of the fourth criterion is to count only cross-links that span a long range on the primary structure.

### Identifying linear peptides with modifications (Figure 1D,E)

The identification of modifications formed by formaldehyde on linear peptides was based only on matching the mass of the precursor ion (i.e. MS^1^) to the theoretical mass of the peptide-modification. This approach was taken because of the poor knowledge of where these modifications occur, or how they affect the MS/MS fragmentation. To make the identification more stringent, we set a very narrow tolerance of 1 ppm on the match between the measured and theoretical mass of the peptide plus the modification. Of note, with such a narrow tolerance we did not find any ambiguous cases in which the measured mass could be assigned to more than one peptide. We ran the analysis eight times, each time searching for a different modification: 0.0 (no modification), 12.0, 24.0, 36.0, 48.0, 60.0, 57.0215 (off-target alkylation), and 15.9949 (oxidation) Da. The estimate of the relative abundance of each modification was calculated as the ratio between the number of identified peptides with that modification and the number of identified peptides without modification (0.0 Da). Other search parameters were: Sequence database – the sequences of BSA, Ovotransferrin, and α-Amylase; Protease – trypsin, allowing up to three miscleavage sites; Fixed modification of cysteine by iodoacetamide. Methionine oxidation was not considered.

### Formaldehyde cross-link identification in a small set of proteins (Figure 2, Table S1)

This analysis application exhaustively enumerates all the possible peptide pairs, and compare them to the measured MS/MS events in search of matches that fulfill the criteria below. The search parameters were as follows: Sequence database – the sequences of BSA, Ovotransferrin, and α-Amylase; Protease – trypsin, allowing up to three miscleavage sites; Fixed modification of cysteine by iodoacetamide; Variable modification of methionine by oxidation; Cross-linking can occur on any residue type; Cross-linker is always cleaved; MS/MS fragments to consider: b-ions, y-ions, B-ions (b-ions plus 12.0 Da), and Y-ions (y-ions plus 12.0 Da); MS^1^ tolerance – 6 ppm; MS^2^ tolerance – 8 ppm; Cross-linker mass – one of three possible masses: 24.0, 25.00335, and 26.0067. The three cross-linker masses were considered in turn in the calculation of the theoretical mass of the two cross-linked peptides. These masses address the incorrect reporting of the mono-isotopic mass (Figure S1).

A cross-link was identified as a match between a measured MS/MS event and a peptide pair if it fulfilled five conditions: 1) The mass of the precursor ion is within the MS^1^ tolerance of the theoretical mass of the linked peptide pair (with either of the three possible cross-link masses); 2) At least four modified MS/MS fragments (B and Y) were identified within the MS^2^ tolerance on each peptide; 3) The fragmentation score of the cross-link (defined as the number of all matching MS/MS fragments divided by the combined length of the two peptides) is 1.0 or higher; 4) The peptides are not overlapping in the protein sequence; 5) There is no other peptide pair or linear peptide that match the data with equal or better fragmentation score.

Given the small size of the sequence database, we estimated the false-detection rate (FDR) in the following way. The analysis of data from the 4% FA experiment was repeated ten times with an erroneous cross-linker mass of 61.0, 62.0, 63.0, … 70.0 Da. This led to fragmentation scores that were much lower than the scores obtained with the correct cross-linker mass. On average, 2 erroneous cross-links had a fragmentation score above 1.0 in each decoy run, whereas runs with the correct cross-linker mass (24.0 Da) identified ~60 cross-links above the 1.0 score. We therefore estimate the FDR to be 2 in 60 cross-links or ~3%.

### Formaldehyde cross-link identification in a large set of proteins (Figure 3, Table S3)

This application relied on the complete cleavage of the FA cross-links in order to separately assign a MS/MS fragmentation score to each peptide. This division allows for a practical run time of *O*(*n*) with suitable preprocessing. The search parameters were as follows: Sequence database – comprising the 1692 human proteins that were identified in the samples. Note that runs on the full human proteome (20,000 proteins) are possible, but take up to four hours; Protease – trypsin, allowing up to two miscleavage sites; Fixed modification of cysteine by iodoacetamide; Cross-linking can occur on any residue type; Cross-linker is always cleaved; MS/MS fragments to consider: b-ions, y-ions, B-ions (b-ions plus 12.0 Da), and Y-ions (y-ions plus 12.0 Da); MS^1^ tolerance – 4.2 ppm; MS^2^ tolerance – 6.5 ppm; Cross-linker mass – one of five possible masses: 24.0, 25.00335, 26.0067, 12.0, and 13.00335 Da. All of these masses were considered in turn in the calculation of the theoretical mass of the two cross-linked peptides. The five masses address the incorrect reporting of the mono-isotopic mass (Figure S1), as well as the much less frequent 12 Da reaction.

A cross-link was reported if it fulfilled four conditions: 1) The mass of the precursor ion is within the MS^1^ tolerance of the theoretical mass of the cross-linked peptide pair (with any of the five cross-link masses); 2) Each peptide had at least 19 MS/MS fragments (b, y, B and Y) within the MS^2^ tolerance, OR its fragmentation score (defined as the number of matching MS/MS fragments divided by its length) was 1.8 or higher; 3) The peptides are not overlapping in the protein sequence; 4) There is no other peptide pair or linear peptide that match the data with equal or better fragmentation score.

To estimate the FDR of the reported list of cross-links, we spiked the sequence database with a decoy set comprising some of the sequences in reverse. The proteins used for the decoys were chosen randomly and their number is user defined. In the case of the PC9 lysate, the number of decoy sequences was set to 1/15 the total number of sequences. We therefore estimate the number of false positives in the cross-link list to be 15 times the number of cross-links that include a reverse decoy peptide.

Both analysis applications (small and large sets of proteins) are available for academic use at http://biolchem.huji.ac.il/nirka/software.html.

---

**Figure S1.**
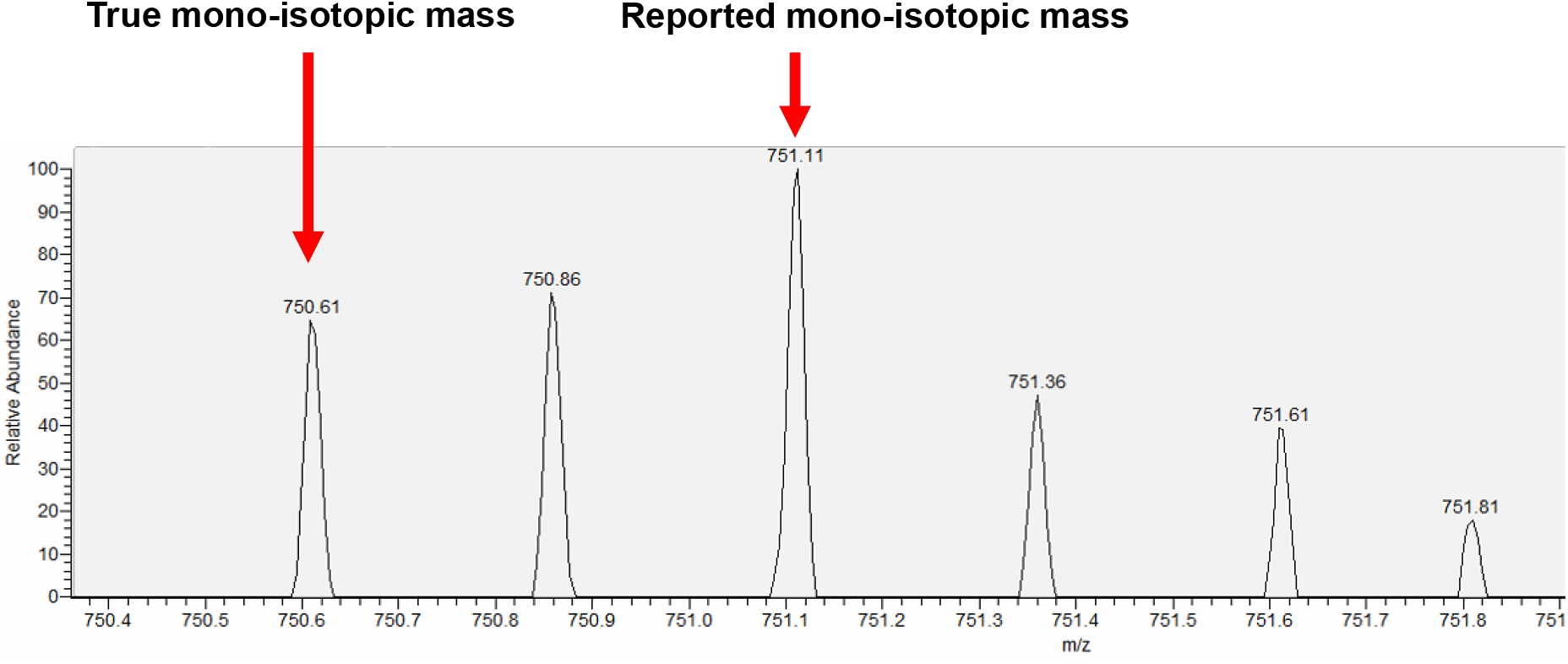
The most intense peak in the isotope series of cross-linked peptides is usually not the mono-isotopic mass. Consequently, the mass-spectrometer often incorrectly report a mass that is approximately 1, 2, or even 3 Da heavier than the true mass (these values being multiples of the mass difference between Carbon-13 and Carbon-12). In Q-Exactive Plus mass spectrometers, which were used in this work, this incorrect assignment of the mono-isotopic mass occurs for about half of the cross-link identifications. The example shown here is an ion of identified cross-link for which the reported mono-isotopic mass was 2 Da heavier than the true mass. This is the cause for the peak broadening around the 24 Da reaction in Figure 1C. In this work, we address this artifact by expanding the search to matches with either the reported mass, the mass minus 1.003 Da, or the mass minus 2.006 Da.

**Figure S2.**
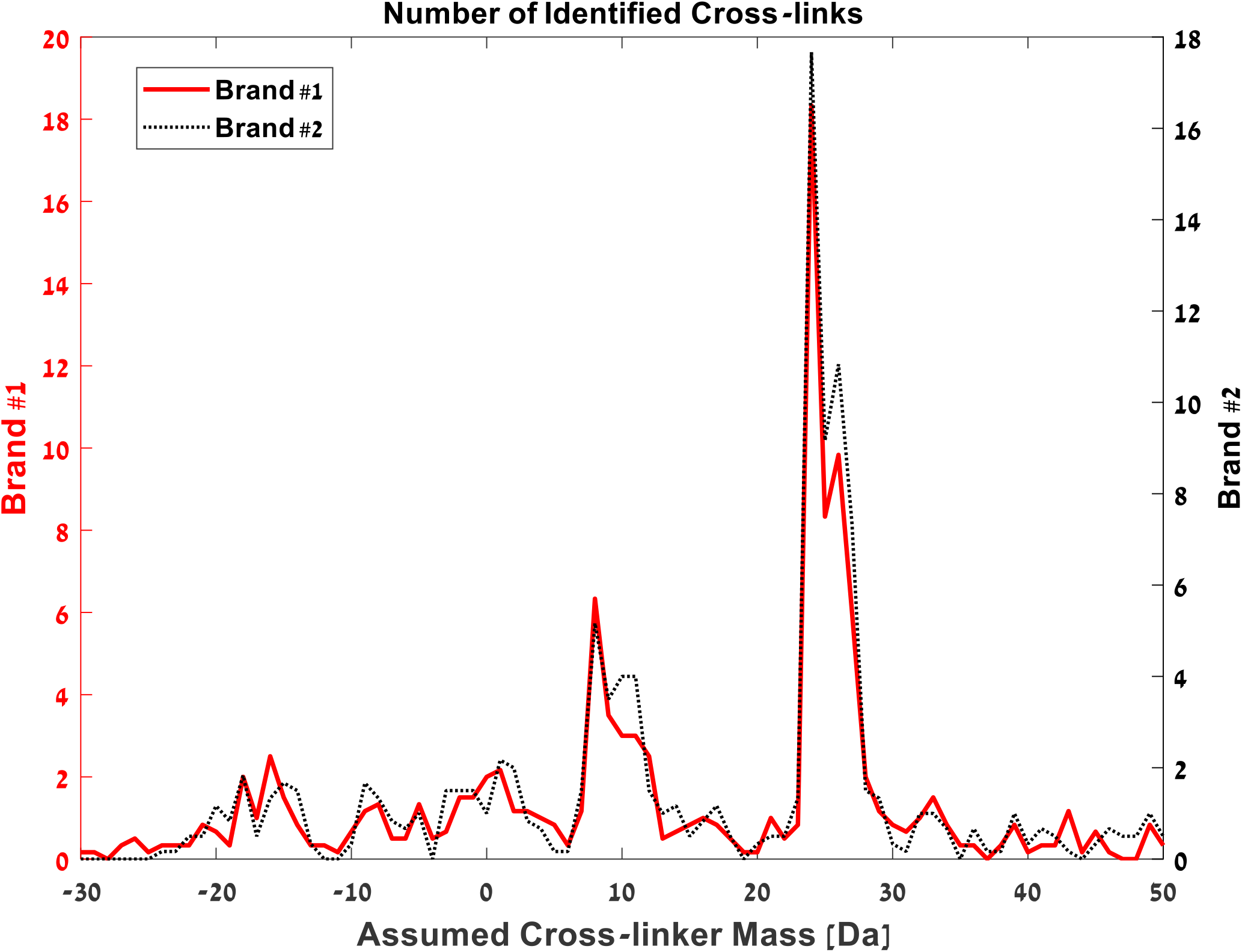
The mass scans described in Figure 1C for data from cross-linking experiments that used two different brands of formaldehyde. Both brands gave essentially the same profiles. The formaldehyde concentration was 1.3%. Brand #1 is from Sigma (product number F8775). Brand #2 is from DAEJUNG chemicals (product number 4044-4400).

**Figure S3.**
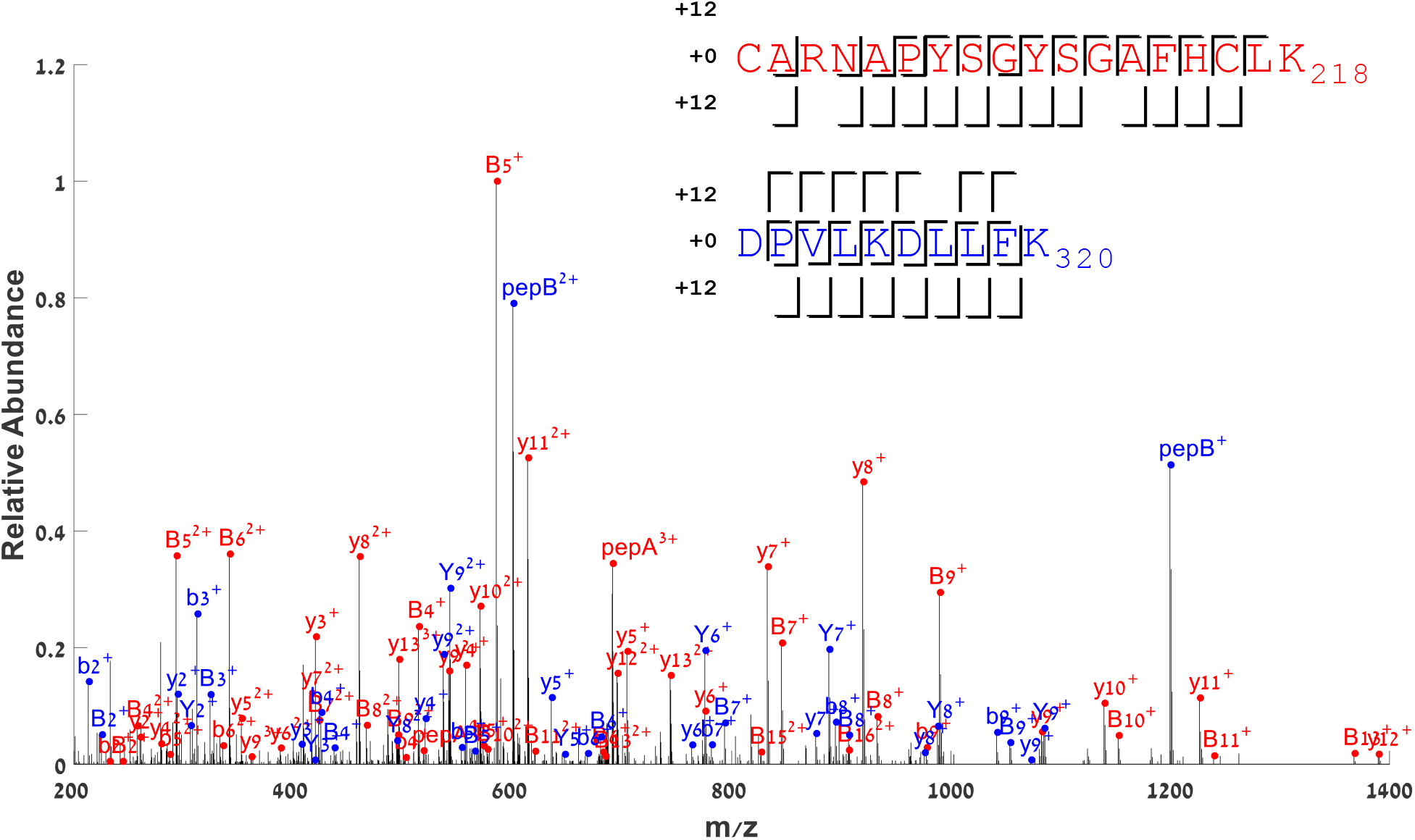
An annotated MS/MS spectrum from HCD fragmentation of two cross-linked peptides from ovotransferrin. PepA and PepB are peaks matching the total mass of the corresponding peptides plus 12 Da. Peaks annotated with capital B or Y match the mass of the corresponding b or y-fragments plus 12 Da.

**Figure S4.**
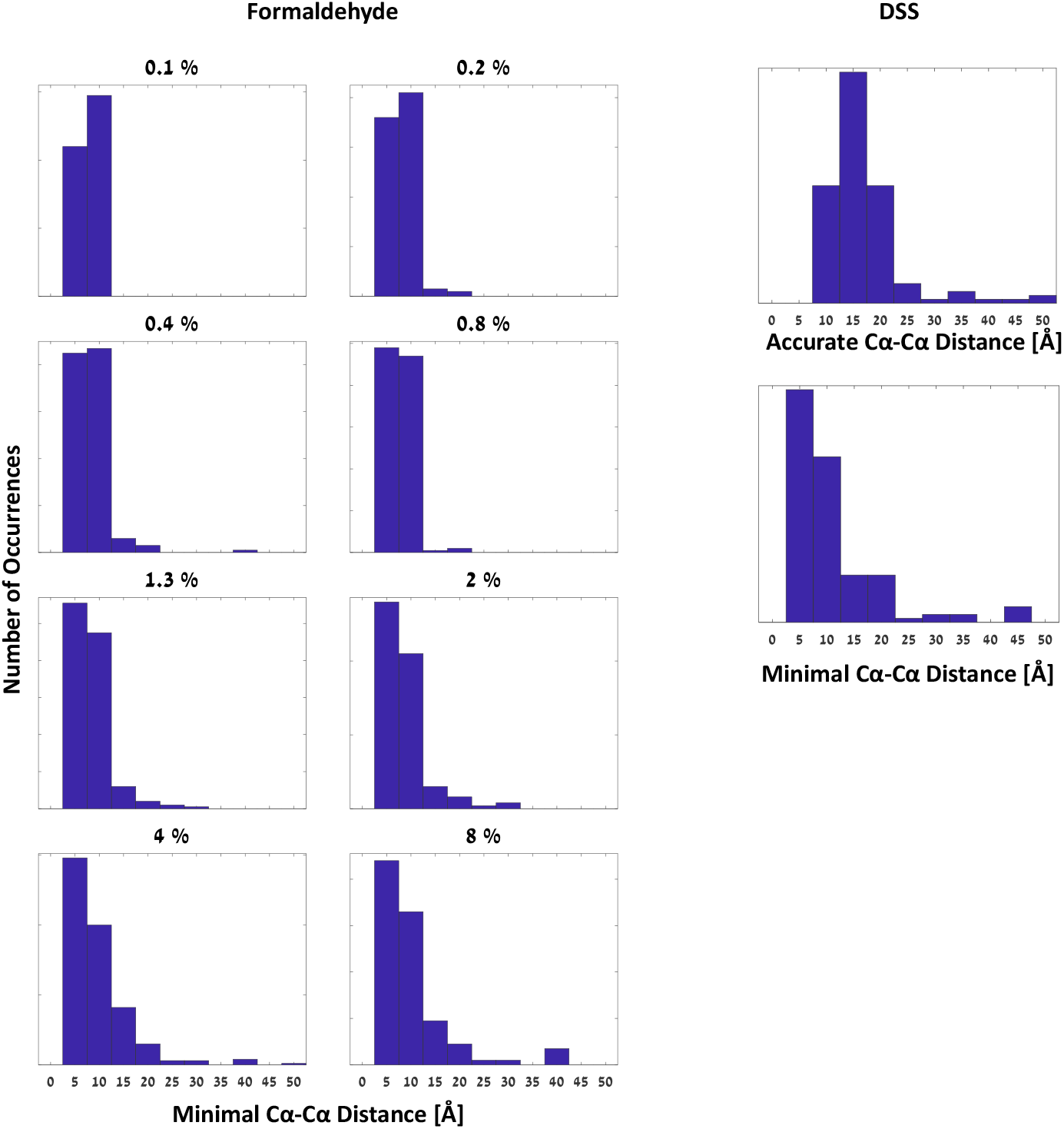
The histogram of the distances between cross-linked peptides identified in the eight formaldehyde concentrations **(left)** and cross-linking with 3mM DSS (disuccinimidyl suberate) **(right)**. The cross-link distance was estimated as the minimal Cα-Cα distance found between the peptide pair on the crystallographic structure. For DSS cross-linking the linked sites are known without ambiguity and accurate Cα-Cα distances can be calculated **(right, top)**.

