## Supplemental Figure 5 for "Mass spectrometry reveals the chemistry of formaldehyde cross-linking in structured proteins"

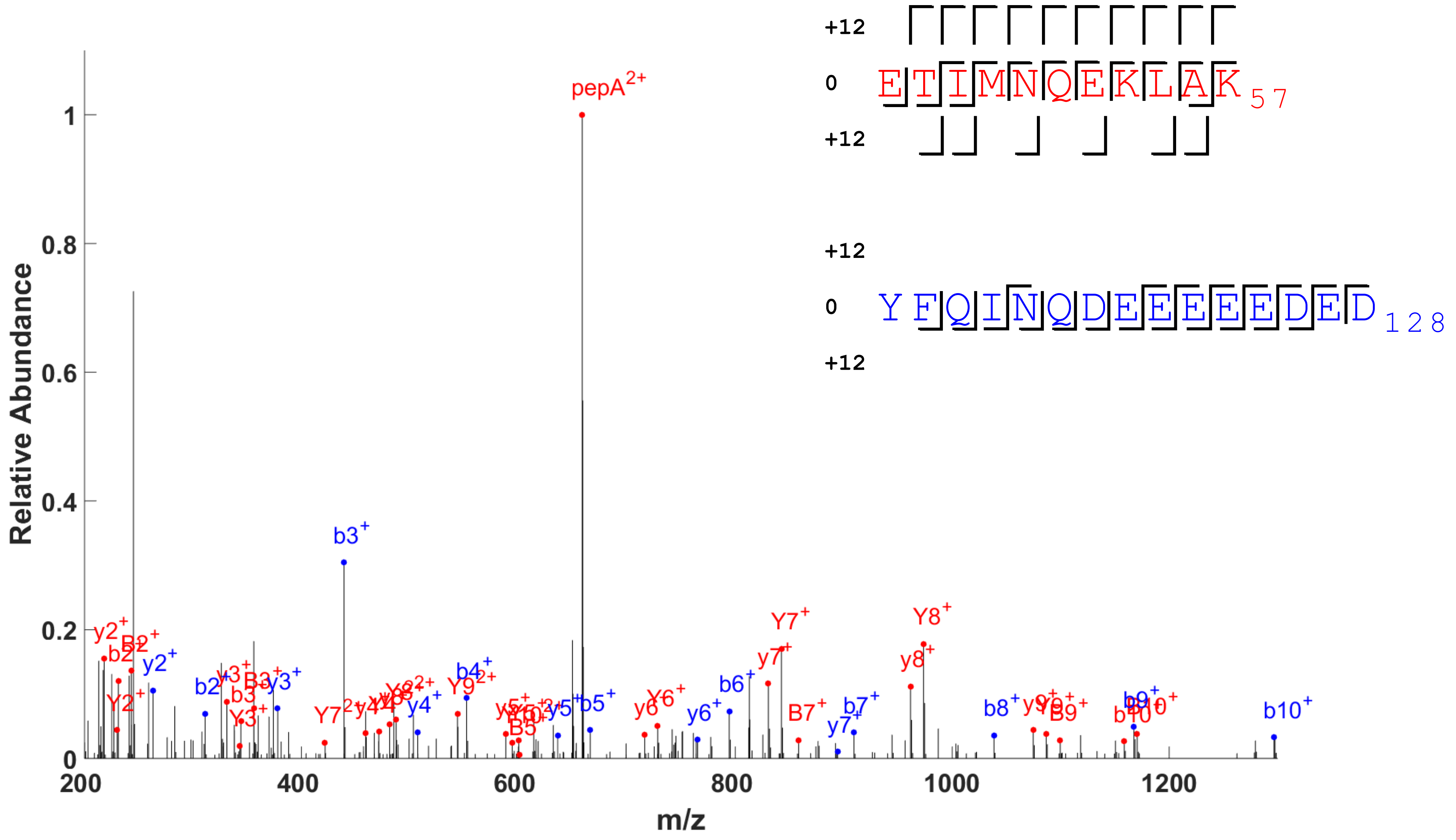


```
+12 A - 1315.68057

12  :  bb b b bb
 0  : bbb      b
SEQ : ETIMNQEKLAK
 0  :   yyyyyyyyy
12  :  yyyyyyyyyy


12  :
 0  :  bbbbbbbbbbbb
SEQ : YFQINQDEEEEEDED
 0  :      y    yyyyyyy
12  :      |
```

BTF3_HUMAN RL22_HUMAN 47 114 ETIMNQEKLAK YFQINQDEEEEEDED-Cterm mass=12

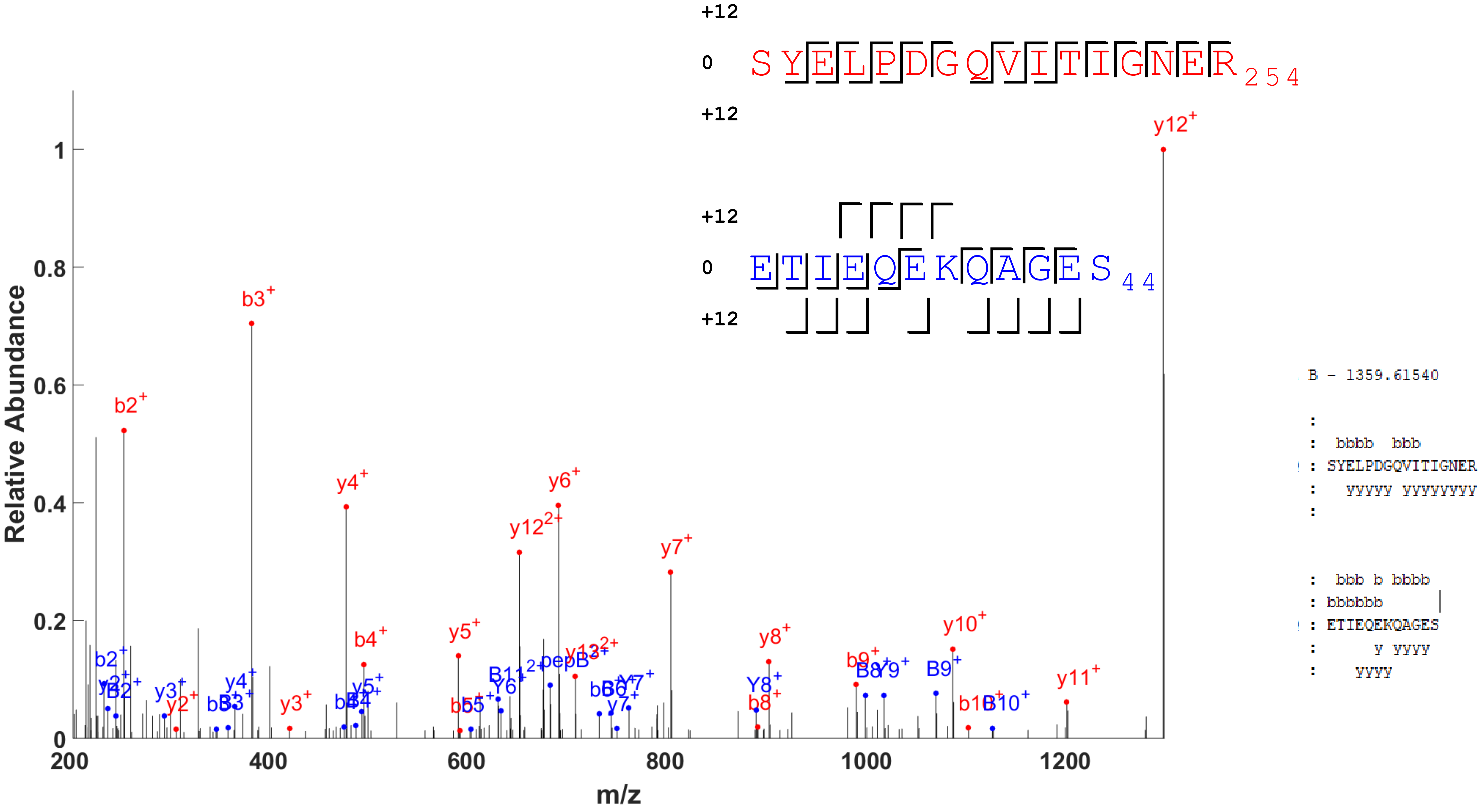


ACTB_HUMAN TYB4_HUMAN 239 33 SYELPDGQVITIGNER ETIEQEKQAGES-Cterm mass=12

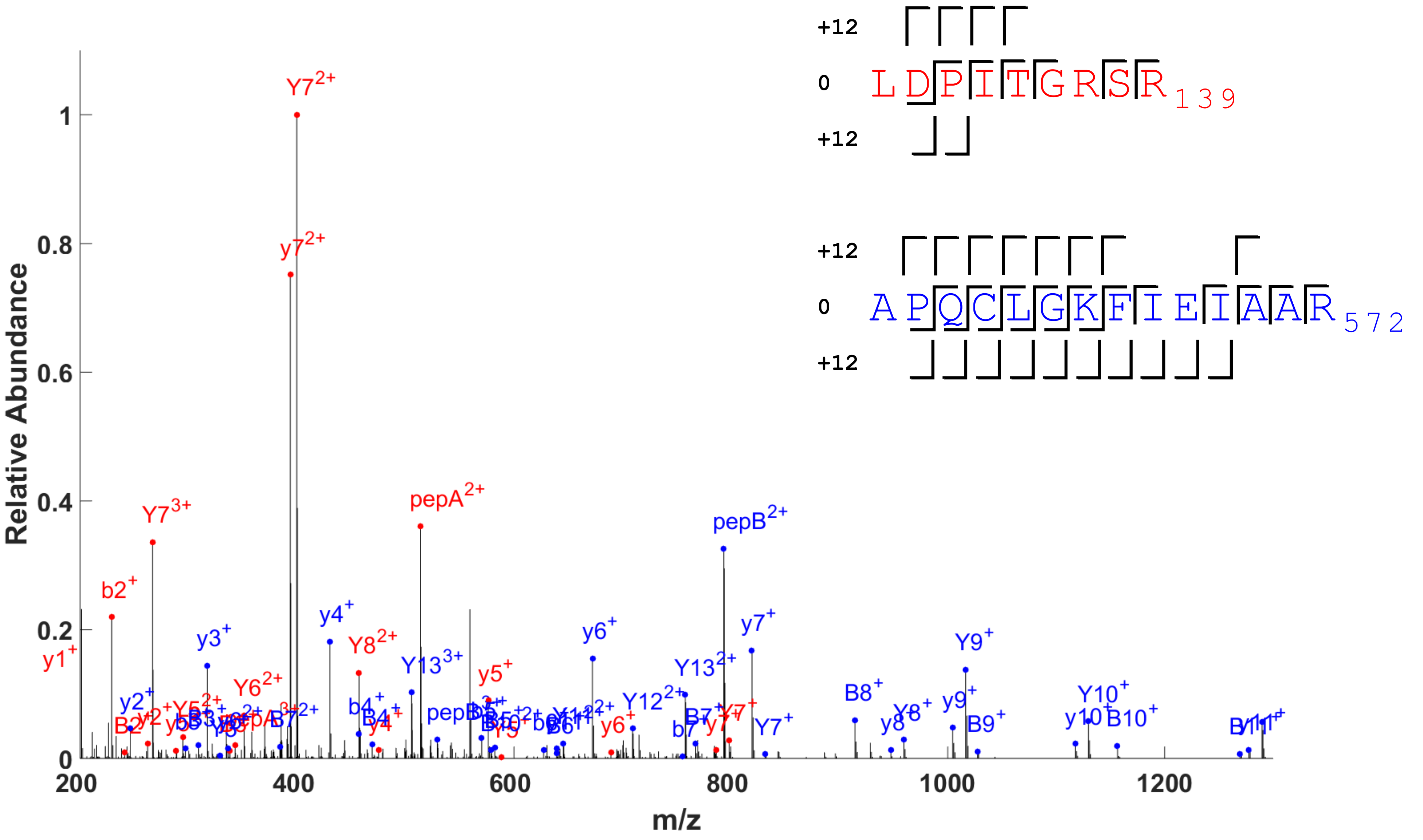


```
+12 A - 1025.56179
+12 B - 1584.84462

12  :  bb
 0  :  b
SEQ : LDPITGRSR
 0  :   yyyy yy
12  :  yyyy

12  :  bbbbbbbbbb
 0  :  bbbbbb
SEQ : APQCLGKFIEIAAR
 0  :   yyyyyyy yyyy
12  :  yyyyyyy   y
```

HNRPD_HUMAN HNRPU_HUMAN 130 559 LDPITGRSR APQCLGKFIEIAAR mass=24

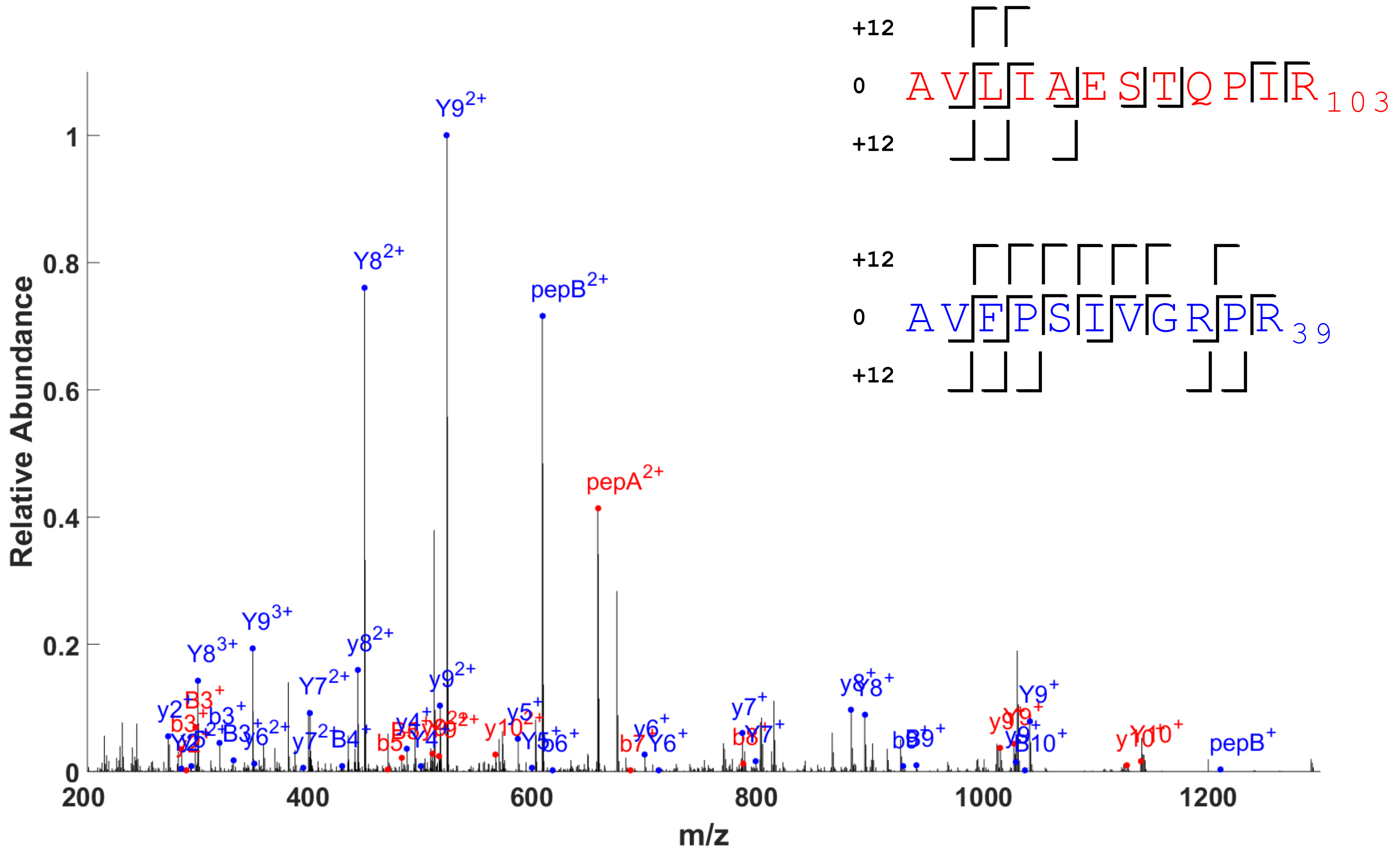


```
+12 A - 1308.74014
+12 B - 1209.69822

12  :  bb b
 0  :  bb b bb
SEQ : AVLIAESTQPIR
 0  :   yy      yy
12  :   yy

12  :  bbb    bb
 0  :  bb  b  b
SEQ : AVFPSIVGRPR
 0  :   yyyyyy yy
12  :   yyyyyy y
```

P210L_HUMAN ACTB_HUMAN 92 29 AVLIAESTQPIR AVFPSIVGRPR mass=24

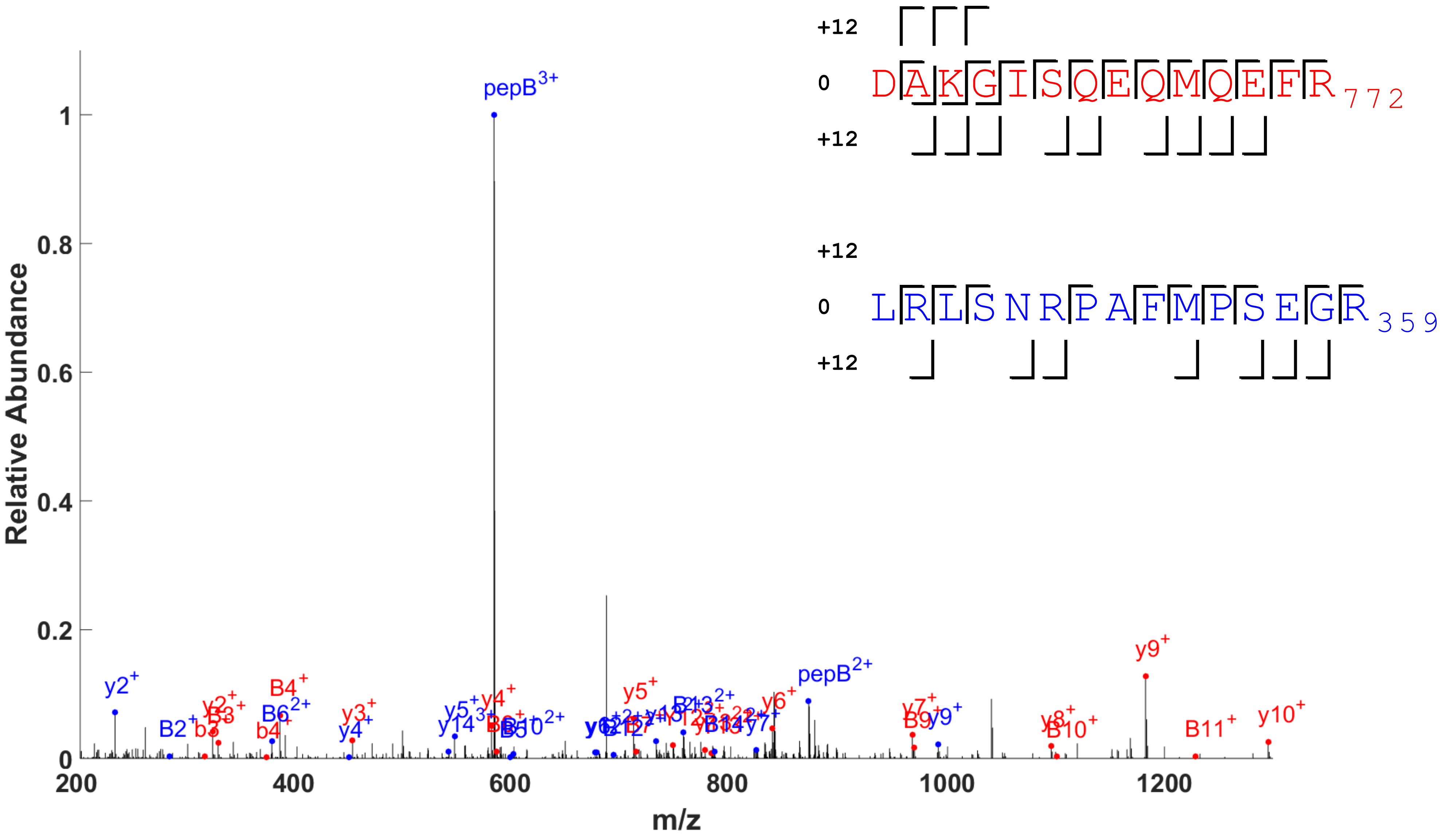


```
+12 B - 1741.90459

 12  :  bbb bb bbbb
  0  :  bbb
SEQ  : DAKGISQEQMQEFR
  0  :  y yyyyyyyyyyy
 12  :  yyy


 12  :  b  bb   b bbb
  0  :
SEQ  : LRLSNRPAFMPSEGR
  0  :  yyy  y yyyy yy
 12  :
```

ACTN4_HUMAN ACTN1_HUMAN 758 345 DAKGISQEQMQEFR LRLSNRPAFMPSEGR mass=24

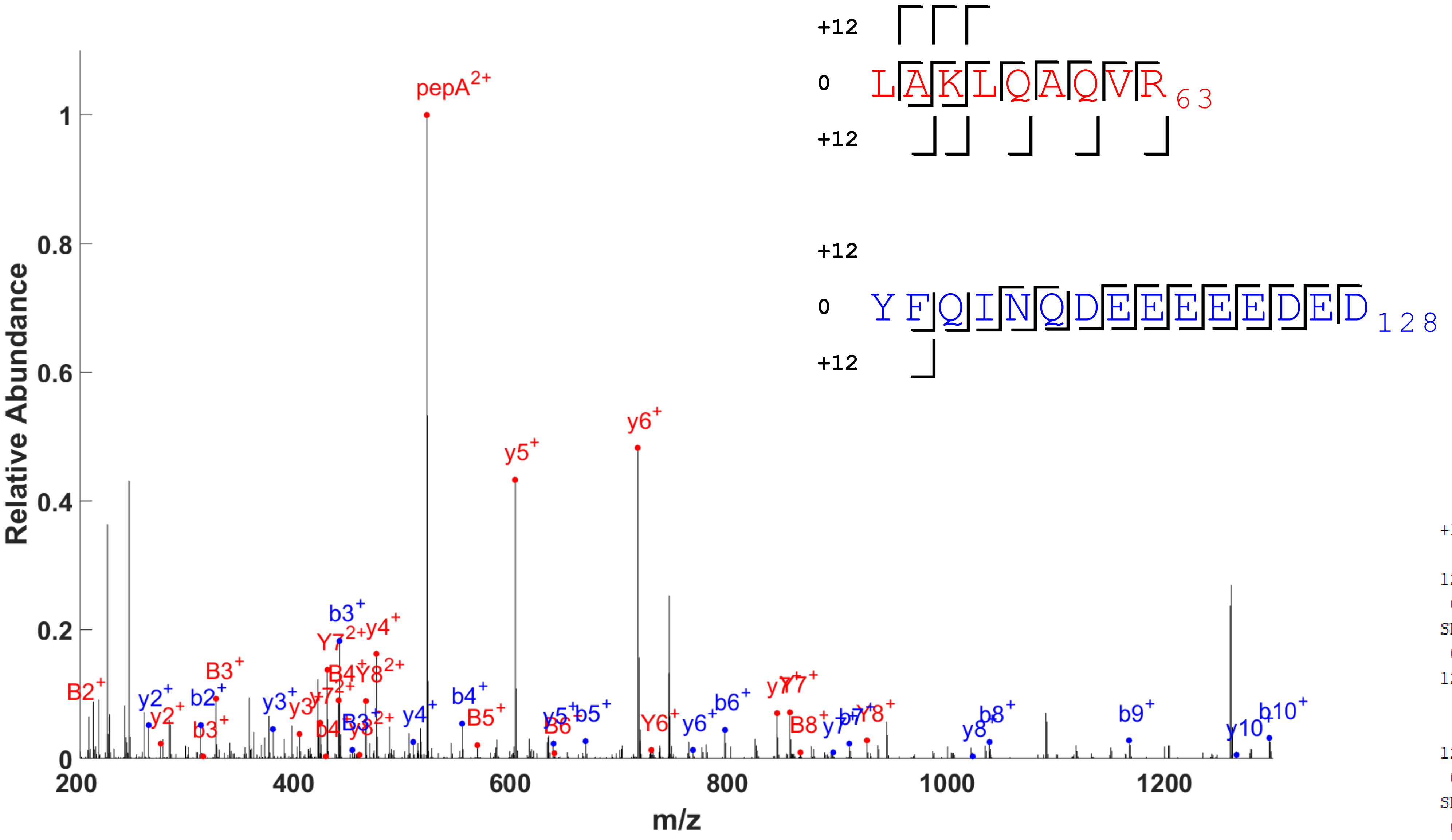


```
+12 A - 1037.63456

12  :  bbbbb b
 0  :  bbb
SEQ : LAKLQAQVR
 0  :  yyyyyyyy
12  :  yyy

12  :   b
 0  :  bbbbbbbbbbbbb
SEQ : YFQINQDEEEEEDED
 0  :     yy yyyyyyyy
12  :
```

BTF3_HUMAN RL22_HUMAN 55 114 LAKLQAQVR YFQINQDEEEEEDED-Cterm mass=12

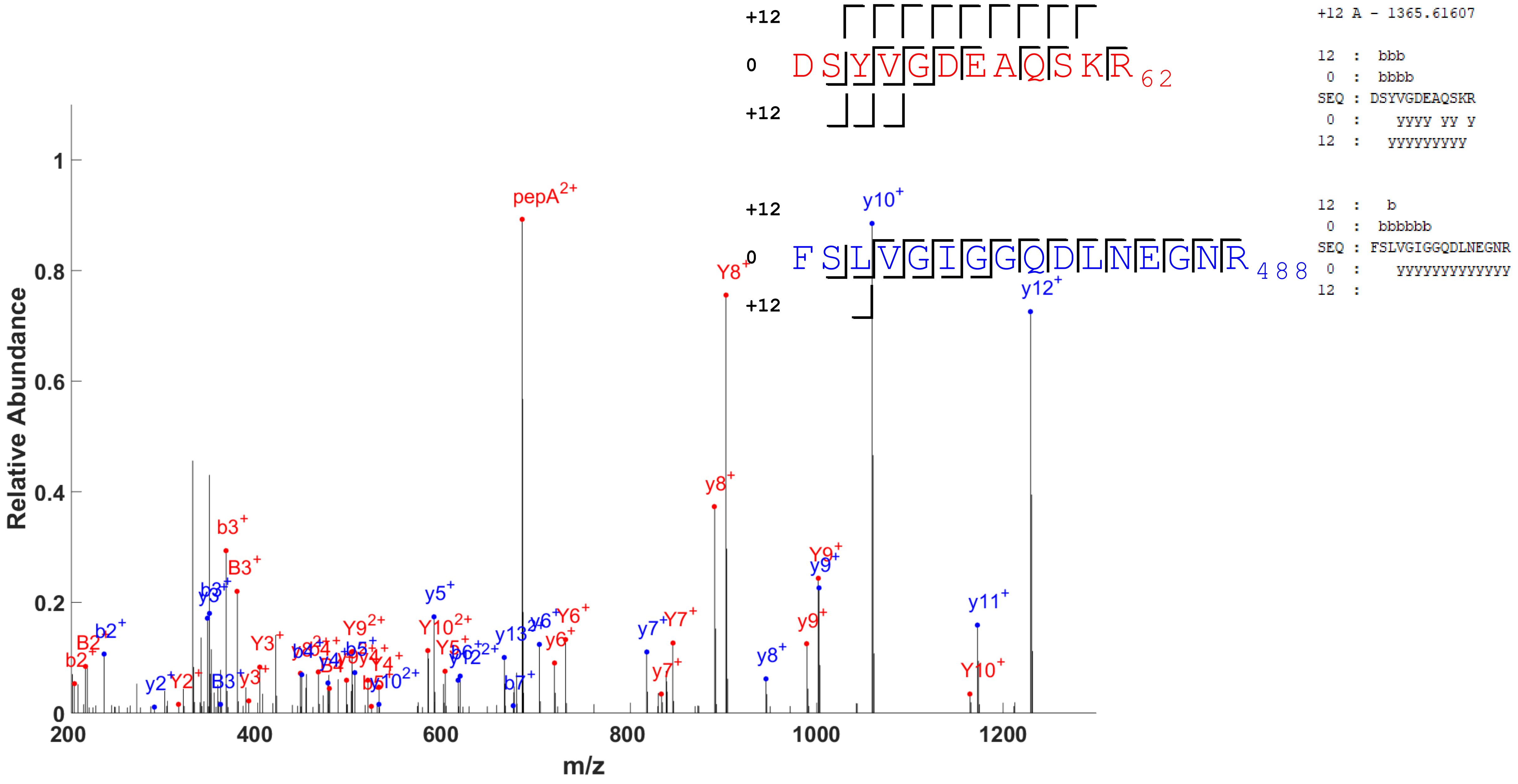


ACTB_HUMAN PLSL_HUMAN 51 473 DSYVGDEAQSKR FSLVGIGGQDLNEGNR mass=12

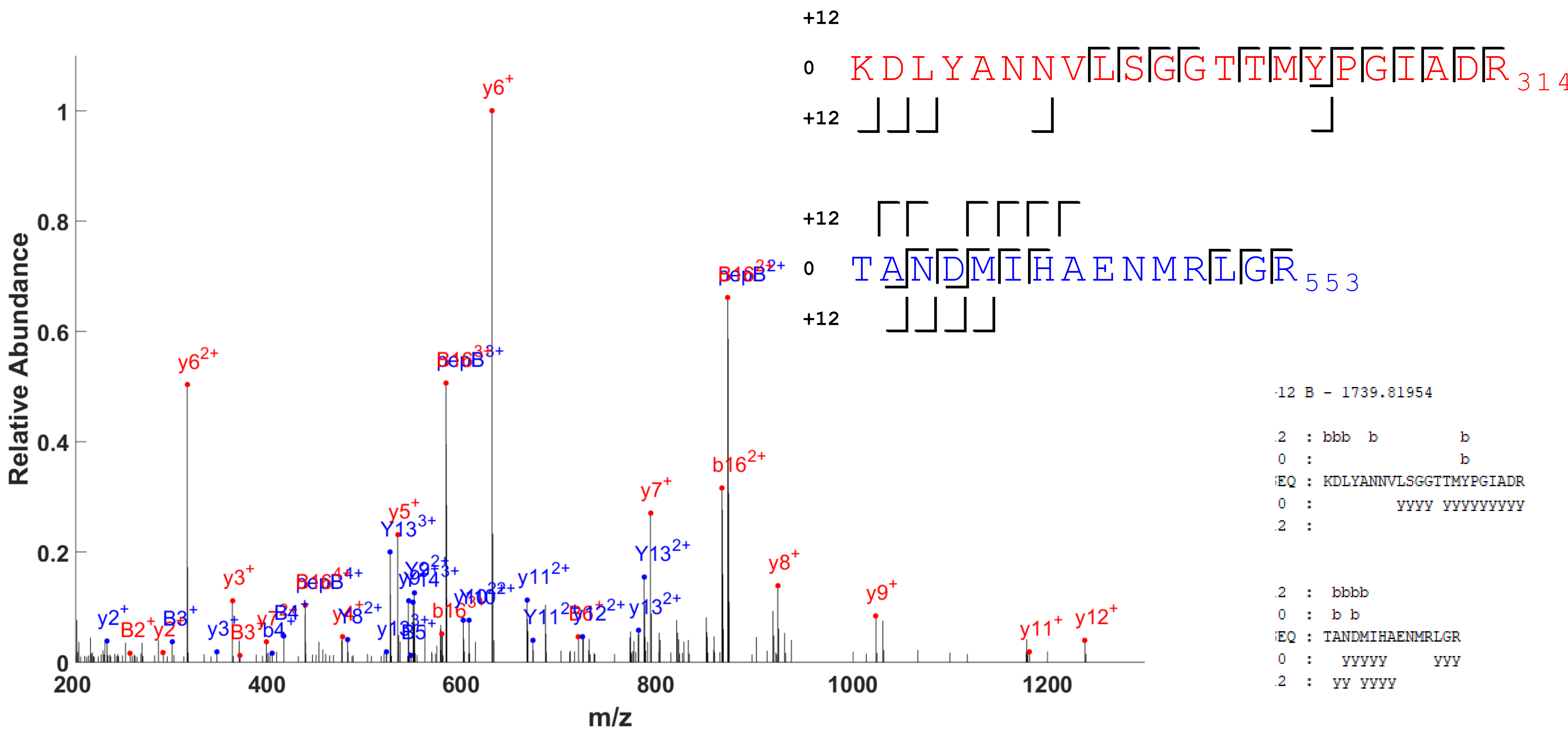


ACTC_HUMAN MOES_HUMAN 293 539 KDLYANNVLSGGTTMYPGIADR TANDMIHAENMRLGR mass=12

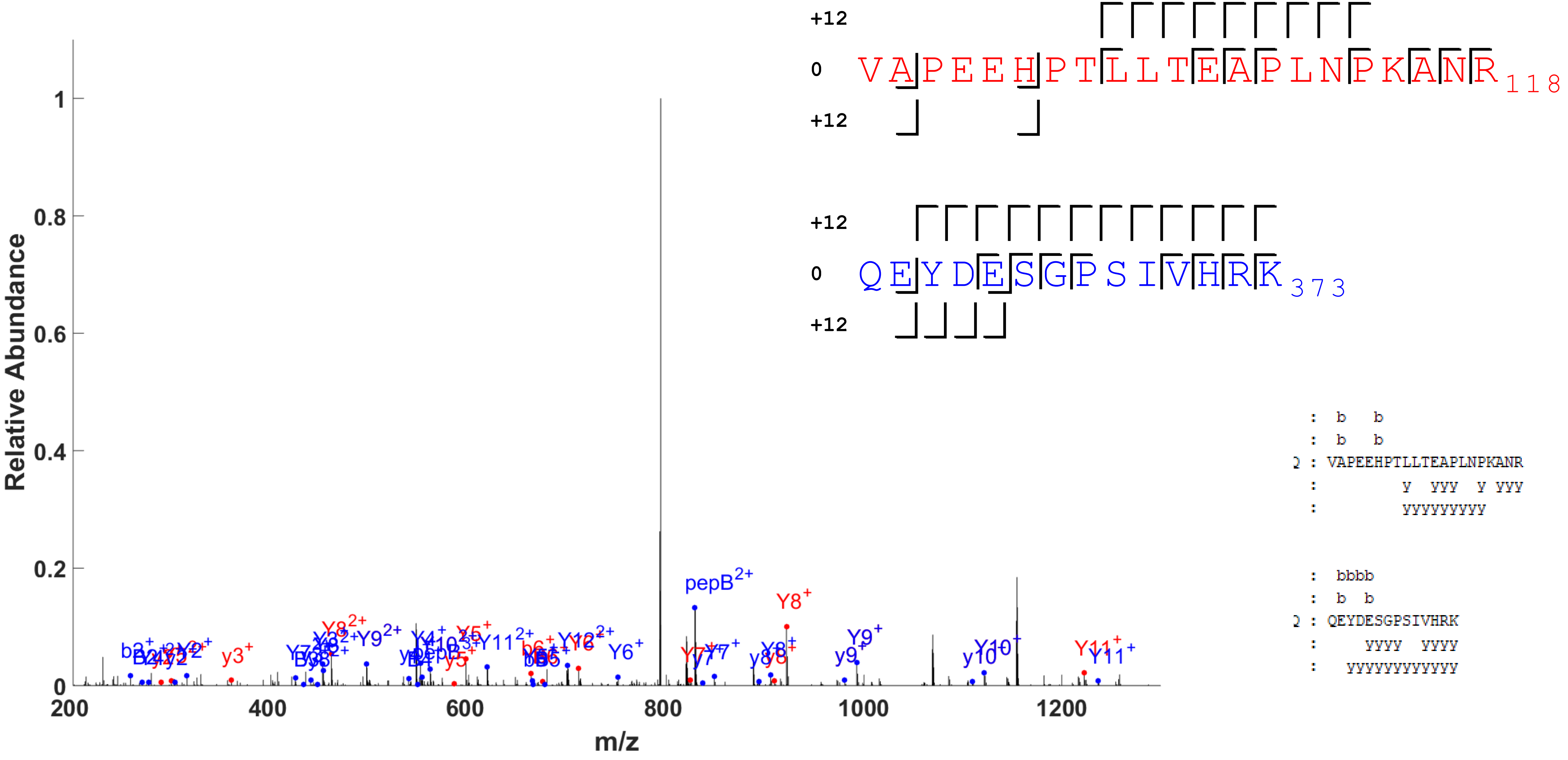


ACTC_HUMAN ACTB_HUMAN 98 360 VAPEEHPTLLTEAPLNPKANR QEYDESGPSIVHRK mass=24

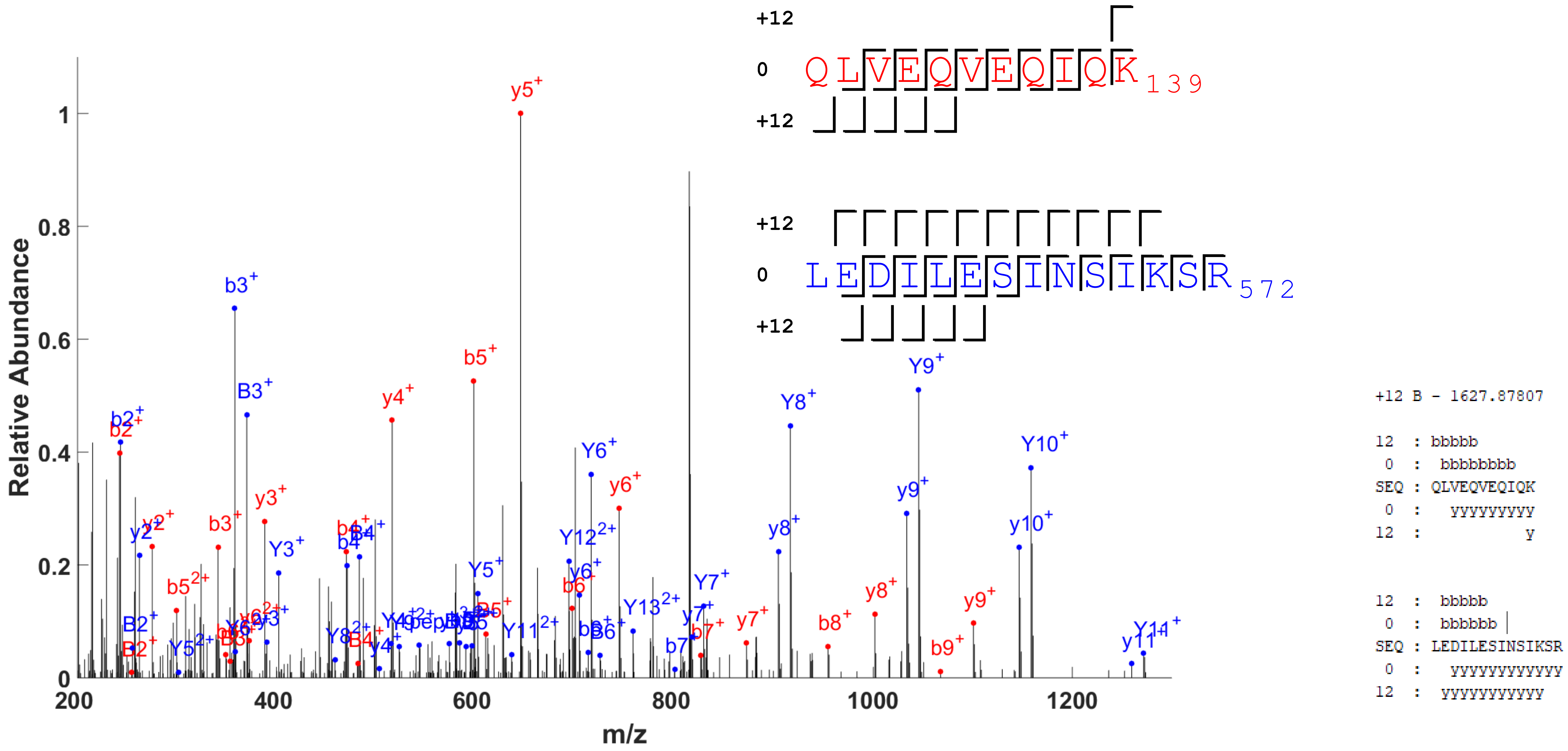


TMED9_HUMAN TMED5_HUMAN 170 153 QLVEQVEQIQK LEDILESINSIKSR mass=12

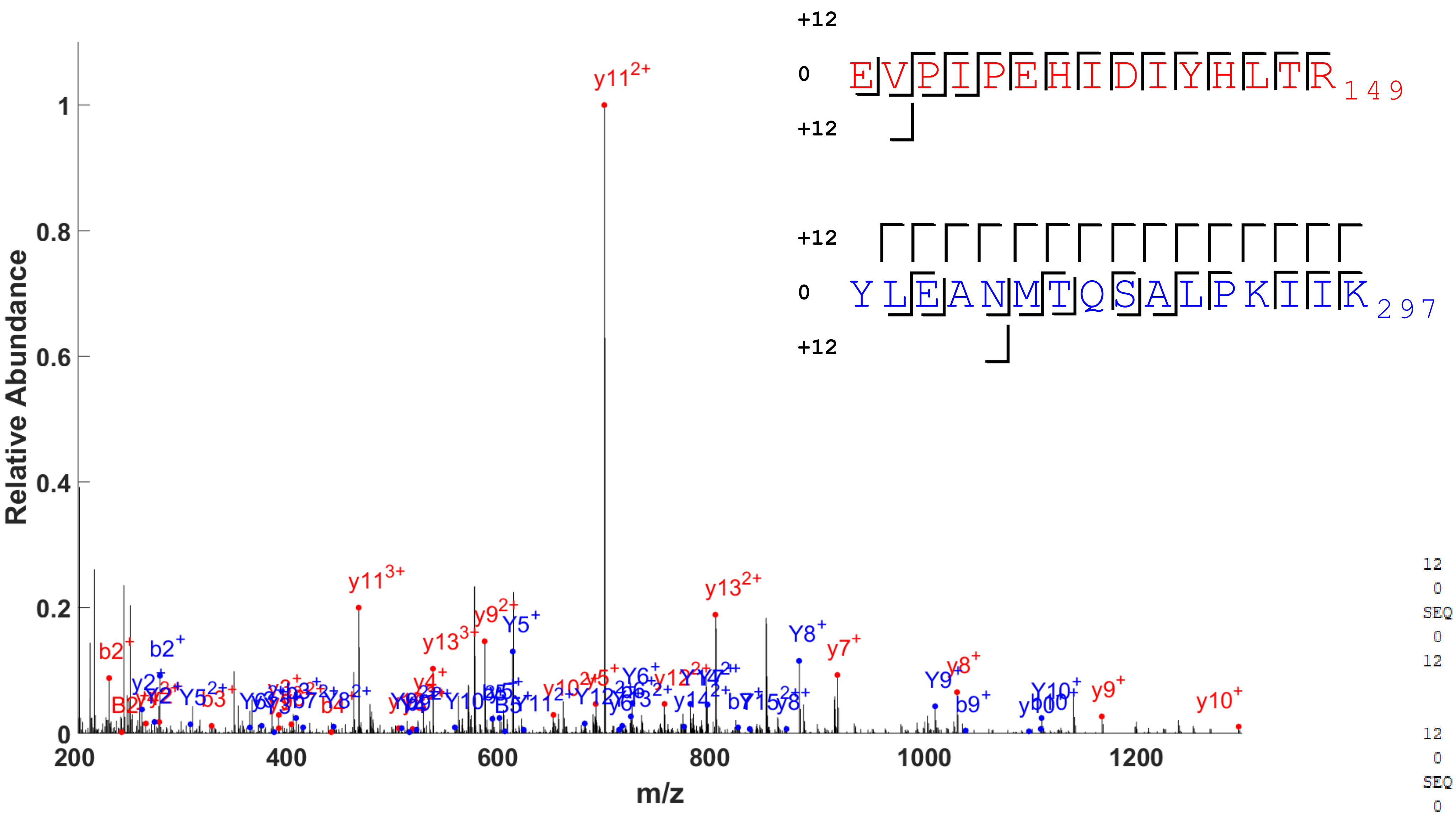


```
12  :  b
 0  : bbbb
SEQ : EVPIPEHIDIYHLTR
 0  :   yyyyyyyyyyyyy
12  :

12  :     b
 0  :  bb bbb bb
SEQ : YLEANMTQSALPKIIK
 0  :    y   y y yy yyy
12  :  yyyyyyyyyyyyyyy
```

ABCF2_HUMAN OLA1_HUMAN 135 282 EVPIPEHIDIYHLTR YLEANMTQSALPKIIK mass=12

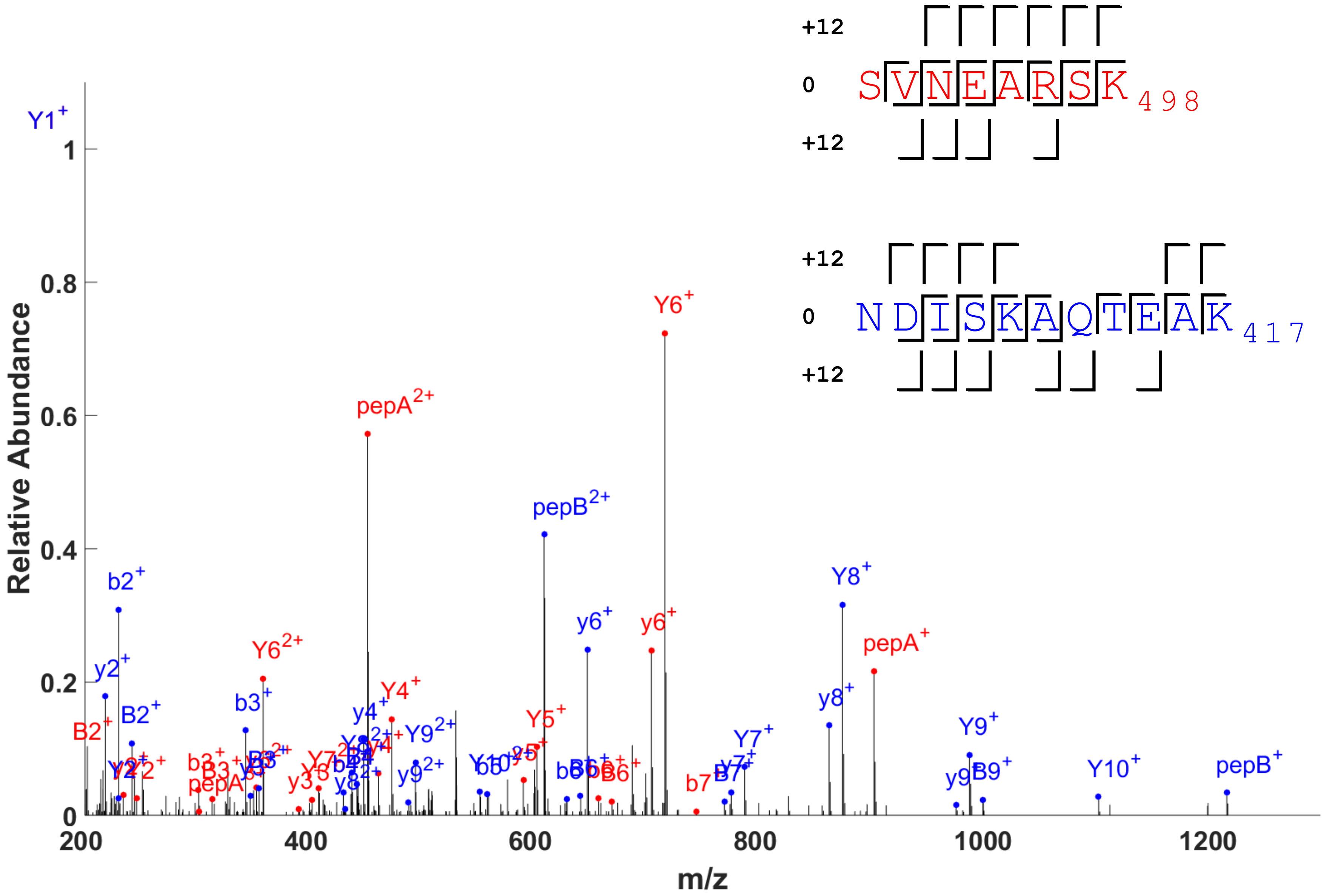


```
+12 A - 901.46174
+12 B - 1215.60953

12  :  bbb b
 0  :  bbb bb
SEQ : SVNEARSK
 0  :   yyyyyy
12  :  yyyyyyy

12  :  bbb bb b
 0  :  bbbbb
SEQ : NDISKAQTEAK
 0  :   yyyy yyyy
12  :  yyyy    yy
```

SMC4_HUMAN SMC2_HUMAN 491 407 SVNEARSK NDISKAQTEAK mass=24

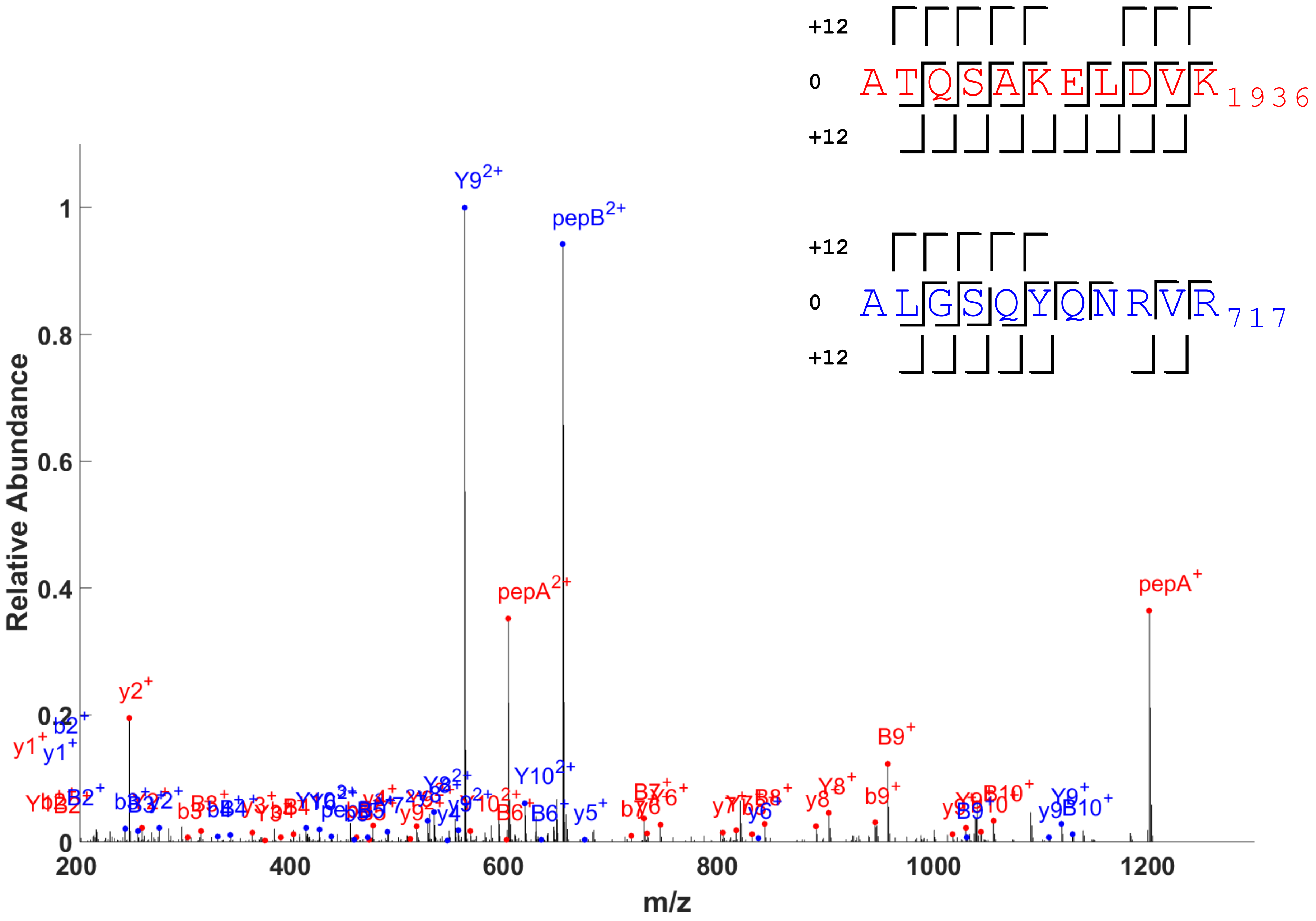


```
+12 A - 1200.63501
+12 B - 1302.67929

12  :  bbbbbbbbb
 0  :  bbbb bbbb
SEQ : ATQSAKELDVK
 0  :   yyyy yyyy
12  :  yyyyy  yyy

12  :  bbbbb  bb
 0  :  bbbb
SEQ : ALGSQYQNRVR
 0  :   yy yyy yy
12  :  yyyyy
```

LAMA3_HUMAN LAMC2_HUMAN 1926 707 ATQSAKELDVK ALGSQYQNRVR mass=24

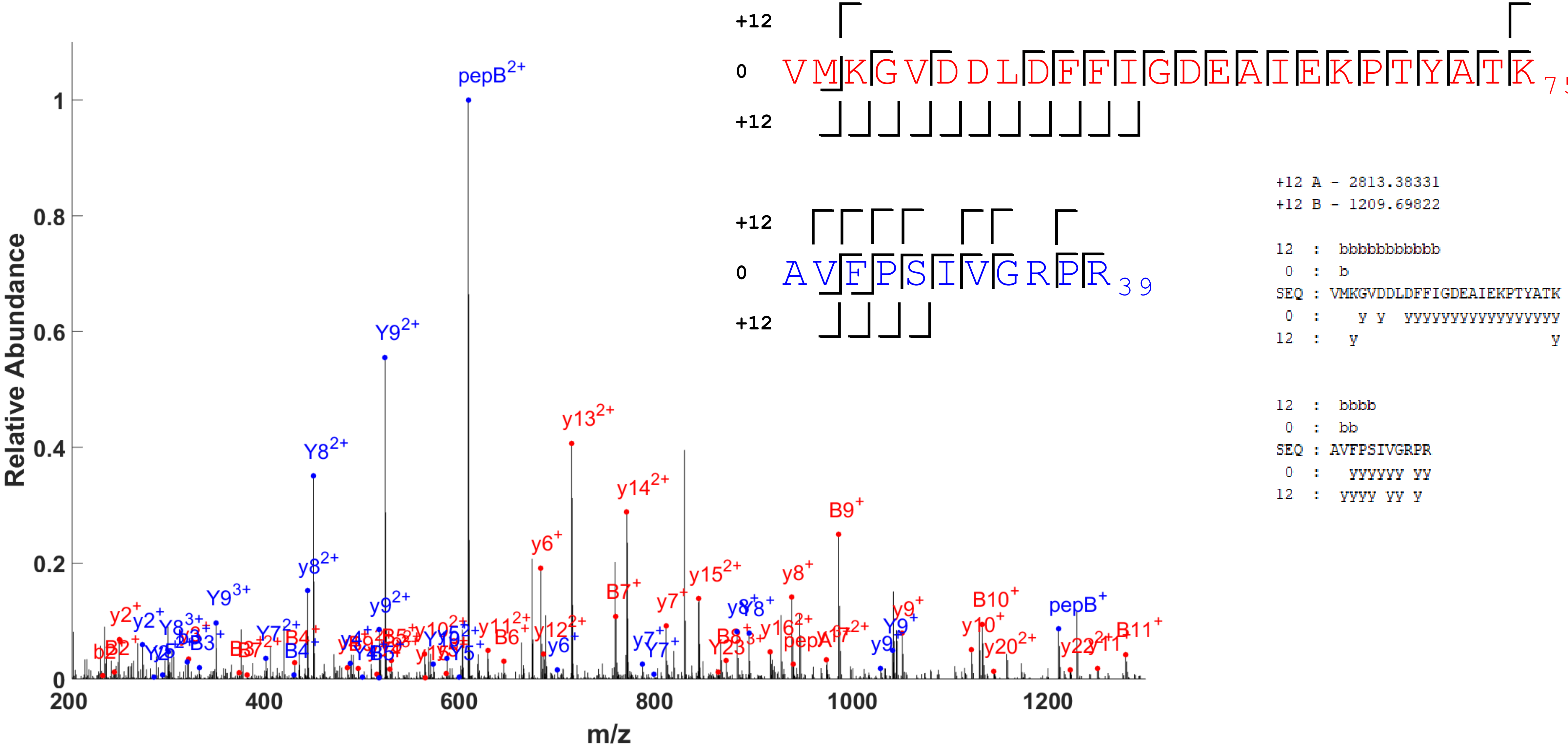


ARP3_HUMAN ACTB_HUMAN 51 29 VMKGVDDLDFFIGDEAIEKPTYATK AVFPSIVGRPR mass=24

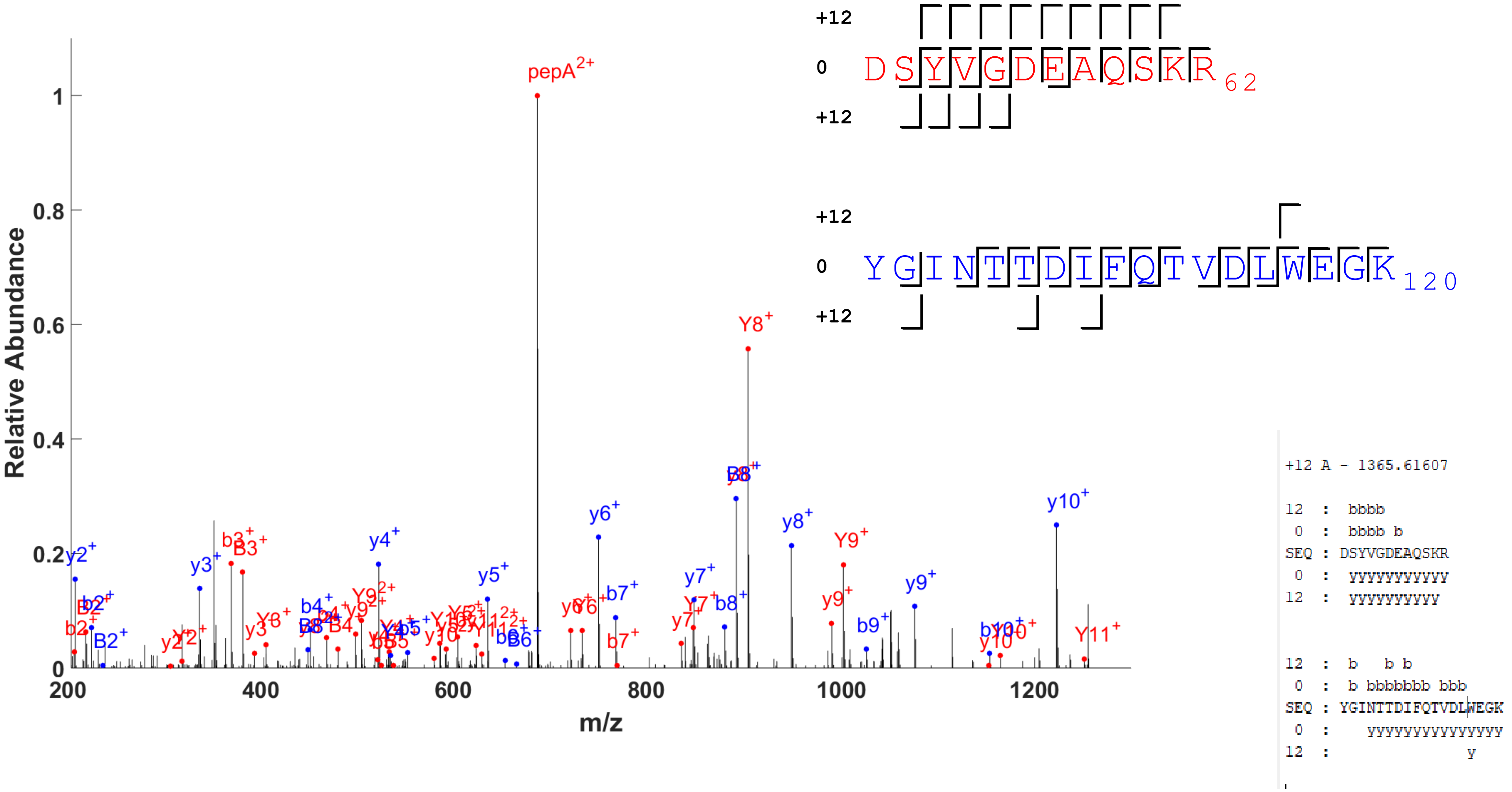


ACTB_HUMAN TAGL2_HUMAN 51 103 DSYVGDEAQSKR YGINTTDIFQTVDLWEGK mass=12

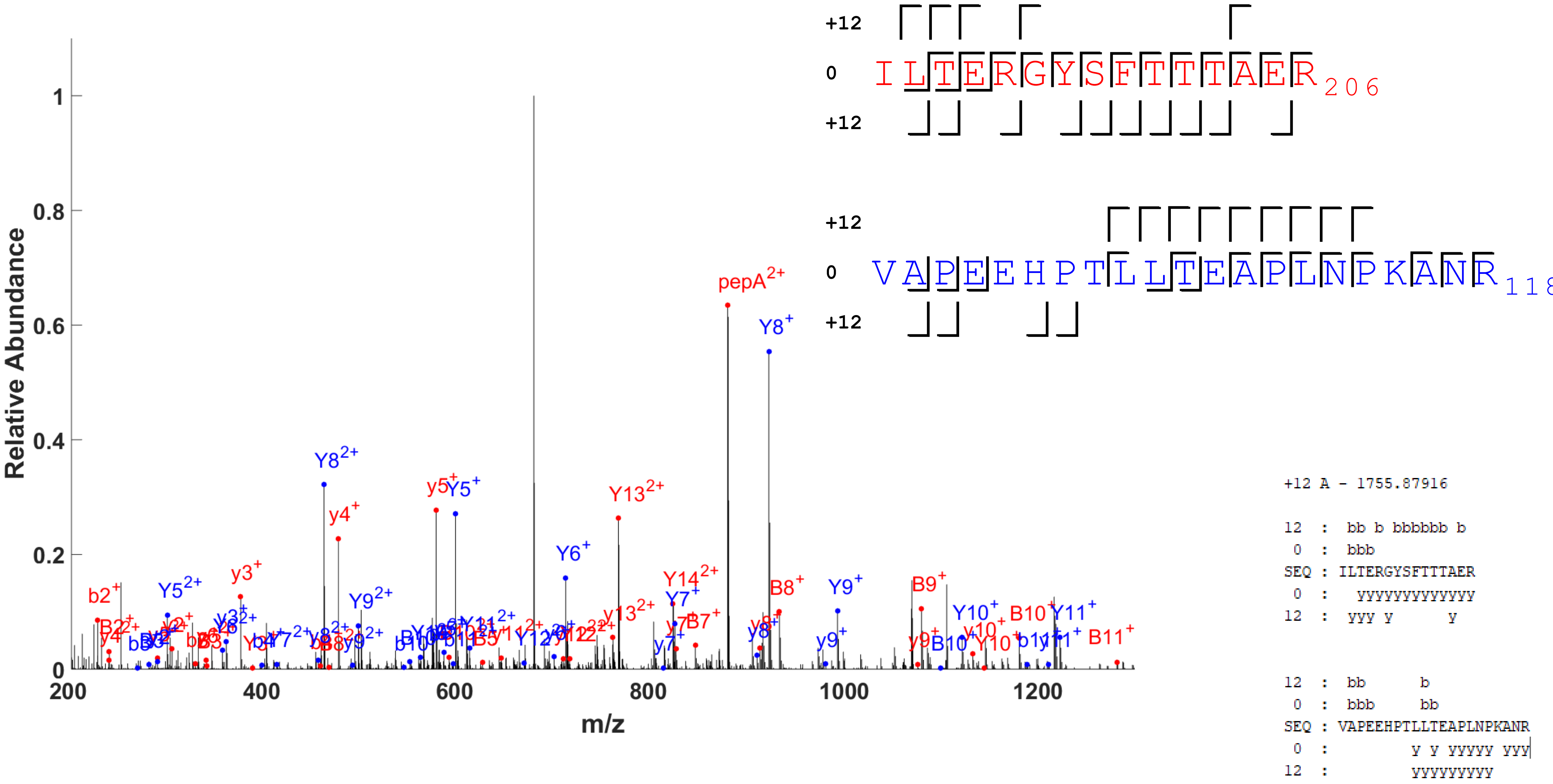


ACTB_HUMAN ACTC_HUMAN 192 98 ILTERGYSFTTTAER VAPEEHPTLLTEAPLNPKANR mass=24
